# Immune context unmasks regulatory effects of Neanderthal and Denisovan introgression

**DOI:** 10.64898/2026.05.29.727880

**Authors:** Zhi Li, Gaspard Kerner, Javier Mendoza-Revilla, Fumitaka Inoue, Simon Fishilevich, Etienne Patin, David Gokhman, Lluis Quintana-Murci, Maxime Rotival

## Abstract

Neanderthal and Denisovan introgression have left a pervasive footprint in the human genome, yet its regulatory consequences remain poorly understood. Here we use a massively parallel reporter assay to characterize the cis-regulatory activity of 4,161 high-frequency introgressed variants across respiratory (A549), hepatic (HepG2), and hematopoietic (K562) cells exposed to immune and infectious stimuli. We find that ~18% of variants show differential activity between archaic and modern alleles, including 94 whose effects are revealed or modulated by stimulation, often in a cell type-specific manner. We identify loci, including *STAT2, IL23A*, and *RNF41*, where clusters of introgressed alleles exert coordinated regulatory effects consistent with adaptive programs. Finally, we dissect the mechanisms underlying the association between Neanderthal introgression and COVID-19 severity and show that the risk allele rs17713054-A, which displays the strongest effect in our assay, increases activity of a TNF-α-responsive enhancer in lung epithelial cells, directly upregulating *SLC6A20*.Together, these findings reveal widespread context-dependent regulatory effects of archaic introgression, with broad evolutionary and biomedical implications.

---

Admixture between modern humans and archaic hominins has resulted in contemporary non-African populations carrying about 2% Neanderthal ancestry, whereas mainland Asians, Native Americans, and Oceanians carry ~0.2-5% Denisovan ancestry^1-5^. In functionally constrained regions, many introgressed segments were rapidly purged by purifying or background selection following introgression, due either to increased selection efficiency in modern humans or to epistatic incompatibilities with human trans-acting factors^6-9^.

Nevertheless, a subset of archaic-inherited variants has persisted and can impact human biology and health^2,9-15^. These alleles, typically segregating at low frequencies, contribute to phenotypic variation primarily through regulatory rather than protein-coding changes, and are depleted from promoters and broadly active enhancers, consistent with selection removing variants with constitutive effects^8,10,11,16^. Accordingly, the effects of introgressed alleles are expected to be highly context-dependent, varying across cell types and environments.

Despite their overall deleterious impact, archaic alleles have also provided a reservoir of adaptive variation^17,18^. Numerous Neanderthal- or Denisovan-inherited haplotypes have reached high frequencies in specific populations, suggesting selective advantage—a process known as adaptive introgression^19^. Such alleles are particularly enriched near genes involved in immune defense, including innate immunity, virus-interacting proteins, and loci of the adaptive immune system (e.g., HLA and T cell receptor genes), indicating that archaic variants may have afforded modern humans increased protection against infections^20-26^.Notably, Neanderthal-inherited haplotypes on chromosomes 3 and 12, present at appreciable frequencies today, have been associated with COVID-19 severity^13,27^. Likewise, archaic variants are enriched in cis-regulatory elements active in immune cells^16,28^ and among immunity-related expression quantitative trait loci (eQTLs)^29-31^.

However, our understanding of which introgressed variants are functional and how they shape the regulatory landscape of human immune responses remains limited. Many archaic variants are geographically restricted, and because most functional studies focus on European populations^32,33^, their regulatory effects are either uncharacterized or difficult to extrapolate due to population differences in linkage disequilibrium (LD). Studies of immune-cell eQTLs have further shown that many regulatory effects manifests only upon stimulation^29-31,34-37^, indicating that a substantial fraction of regulatory variation remain cryptic at baseline. In addition, statistical eQTL fine-mapping often fails to identify the causal regulatory variant among linked archaic and modern alleles, leaving the functional consequences of introgressed variation unresolved. In this context, massively parallel reporter assays (MPRA) provide a powerful alternative to quantify the regulatory effects of thousands of candidate sequences in controlled cellular settings^38-40^, enabling the identification of causal variants, rare or population-specific alleles, and tissue- and context-specific activity^39,41-50^. Yet, previous MPRA studies of archaic-derived variants have focused on basal expression^43-46^, leaving the context-dependent effects of adaptive introgression on immune function unexplored.

Here we characterized, across multiple cell types and stimulation conditions, the cis-regulatory effects of a curated set of 4,161 variants at immunity-related loci with evidence of adaptive introgression across diverse populations. Using a lentivirus-based MPRA (lentiMPRA)^40^, we profiled their regulatory activity at baseline and following exposure to immune cytokines and live pathogens. This approach revealed widespread context-dependent regulatory effects of introgressed haplotypes, providing a functional atlas of how archaic-derived alleles shape human immune responses and infectious disease susceptibility.

## Results

### Prioritizing archaic-introgressed immune variants using lentiMPRA

To investigate the cis-regulatory effects of archaic introgression in immune contexts, we first compiled 183,652 high-confidence archaic single-nucleotide variants (SNV) from eight populations across Europe, Asia, and the Pacific, encompassing diverse evolutionary histories and levels of archaic ancestry (Fig. 1a). To detect additional introgressed haplotypic variation, we expanded this set to include variants in strong LD (*r*^2^≥0.8) with at least two archaic variants separated by >10 kb, capturing (i) putative archaic alleles absent from sequenced archaic genomes and (ii) alleles predating the modern-archaic split that were lost in modern humans and later reintroduced through admixture^51^. This yielded 487,386 putatively introgressed alleles segregating at >5% frequency in at least one population.

**Fig. 1.**
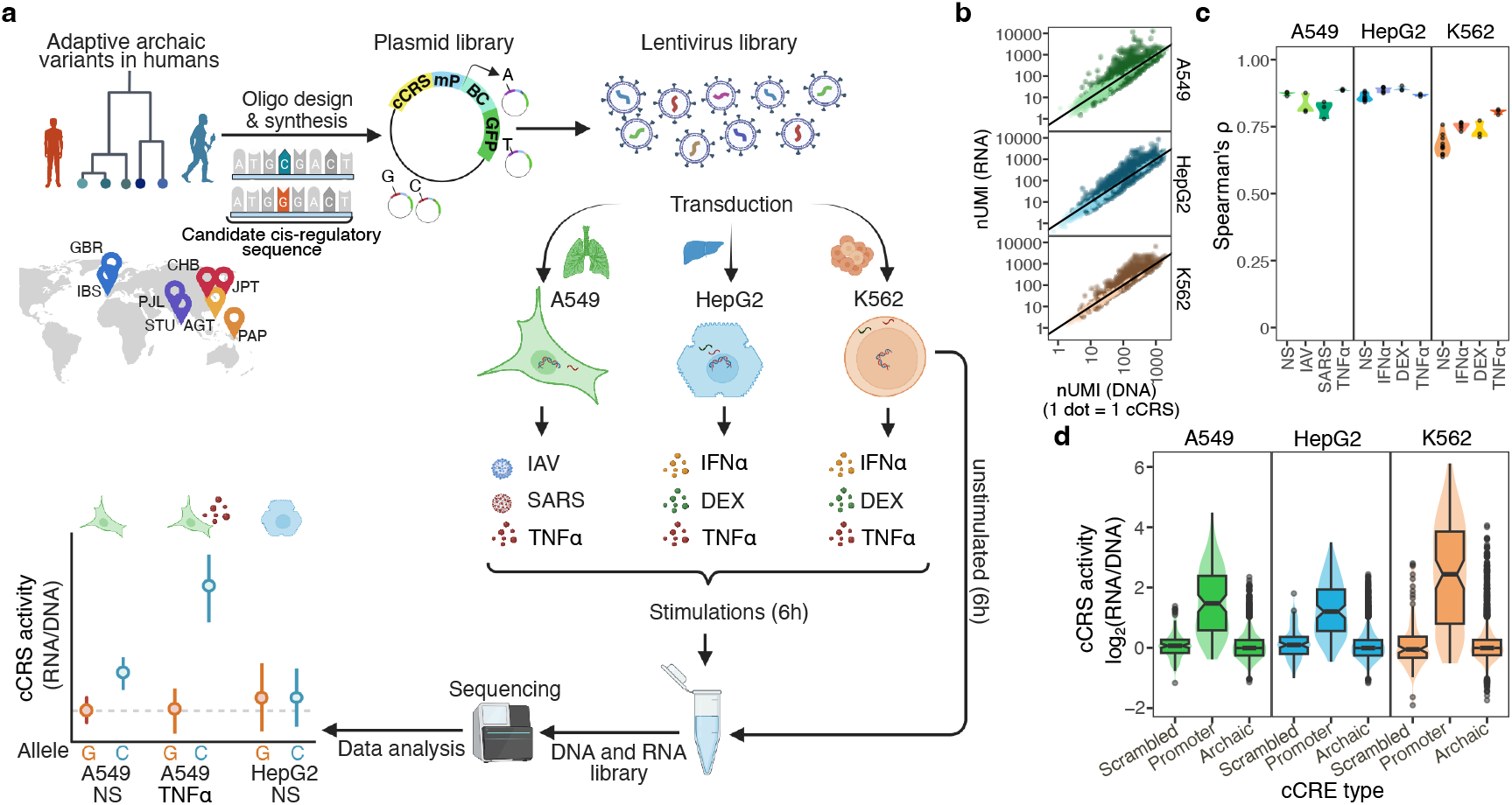
Design and overview of experimental design. **a**, Schematic overview of the lentiMPRA experimental design. A library of 300-nt oligonucleotides was designed to match candidate cis-regulatory elements (cCREs) harboring archaic and modern human alleles at 4,161 introgressed variants, selected from eight populations worldwide. Each sequence was cloned into lentiviral plasmid, upstream of a minimal promoter and unique barcodes, and packaged into lentivirus, followed by transduction into three cell lines: A549, HepG2 and K562. After 6 hours under resting conditions or following stimulation with IFN-α, Dexamethasone (DEX), TNF-α, Influenza A virus (IAV) or SARS-CoV-2 (SARS), DNA and RNA were simultaneously extracted and sequenced to quantify allele-specific transcriptional activity across contexts, enabling the identification of context-specific expression-modulating variants (emVars). **b**, Comparison of barcodes abundance (UMI per million) in RNA and DNA across all three cell types under unstimulated conditions. cCRS with significant regulatory activity (FDR<5%) are highlighted in darker shades. **c**, Pairwise correlations of transcriptional activity profiles between technical replicates within each cell type and condition. **d**, Violin plots showing the distribution of GC-corrected transcriptional activity for random sequences (scrambled controls), promoters (positive controls), and tested introgressed sequences across all three cell types (unstimulated condition).

We next prioritized introgressed variants likely to affect immune regulation by focusing on 2,015 genes involved in innate or adaptive responses and viral infection, selecting introgressed alleles in the top 5% frequency genome-wide (maximum frequency across populations: 0.10-0.93). This identified 4,161 variants located within 10kb of these genes (2,971 variants) or in putative distal enhancers interacting with them (based on promoter-capture Hi-C^52,53^; 1,190 variants) (Extended Data Fig. 1 and Supplementary Table 1). For each variant, we synthesized paired 270-bp genomic sequences centered on the introgressed (archaic) or non-introgressed (modern) allele. Positive controls comprised 113 core promoters of highly expressed genes, while scrambled sequences served as a baseline of activity levels (Methods). In total, 8,548 oligonucleotides (candidate cis-regulatory sequences, cCRS), representing 4,274 candidate cis-regulatory elements (cCREs), were cloned into lentiviral plasmids upstream of a minimal promoter and a unique 15-bp barcode^40^. Each cCRS was linked to a median of >81 barcodes, with >92% associated with >10 barcodes (Supplementary Fig. 1 and Supplementary Table 1). Retaining cCREs with >10 barcodes for both alleles in at least one library resulted in 3,838 introgressed cCREs for downstream analyses. These included variants unambiguously introgressed from Neanderthals (58%, n=2,231) or Denisovans (23%, n=887), as well as variants located on haplotypes shared between both archaic groups (12%, n=442) or not detected in either group (7%, n=278).

To profile regulatory activity across diverse contexts, we transduced the lentiviral library into lung epithelial (A459 adenocarcinoma), hepatic (HepG2 hepatoblastoma), and hematopoietic progenitor (K562 leukemia) cells, selected for their relevance to immunity and infection, and compatibility with lentiMPRA (Supplementary Note 1). Forty-eight hours post-transduction, cells were stimulated for 6 h with interferon-α (IFN-α), dexamethasone (DEX), or tumor necrosis factor-α (TNF-α) in HepG2 and K562 cells, and with influenza A virus (IAV), SARS-CoV-2, or TNF-α in A549 cells (Supplementary Note 2). Each condition included 3-5 replicates, with stimulated cells assayed alongside matched unstimulated controls. Sequencing recovered 79% and 88% of oligo-associated barcodes per DNA and RNA replicate, respectively. Cis-regulatory activity was estimated as the GC-corrected mean RNA/DNA ratio across barcodes (Fig. 1b, Supplementary Note 3 and Supplementary Fig. 2), showing high reproducibility across replicates (Spearman’s ρ=0.62-0.90) that increased with barcode number (Fig. 1c and Extended Data Fig. 2a). Positive controls displayed higher transcriptional activity than scrambled sequences (Δmedian activity >2.2, Wilcoxon rank-sum *P*<10^-77^), whereas introgressed cCREs showed a broad activity distribution, with only a small fraction reaching promoter-like levels (Fig. 1d), consistent with the depletion of introgressed variants from promoters^8^. These results establish a framework for quantifying the context-dependent cis-regulatory effects of archaic introgression.

### Transcriptional activity and regulatory context of introgressed cCREs

To characterize context-specific activity, we performed UMAP on batch-corrected transcriptional profiles, which clustered transduced cells primarily by cell type (33% of variance), followed by stimulation (5%) (Fig. 2a and Supplementary Table 1). Active cCREs were identified using a barcode permutation strategy to detect elements whose RNA/DNA ratios differ significantly from the median of all barcodes (Supplementary Fig. 3). At 5% FDR, 728-2,005 active cCREs were detected per condition, with >90% showing consistent directional effects across alleles (replication *P*<0.01, Spearman’s ρ=0.85-0.93; Extended Data Fig. 2b). Across conditions, 47% (n=1,250) increased transcriptional activity (‘upregulating cCREs’), 49% (n=1,311) decreased it (‘downregulating cCREs’), and 4% (n=114) showed allele- or context-dependent directionality (Fig. 2b and Supplementary Table 1). Most cCREs showed reproducible effects across contexts, with 87% of upregulating and 84% of downregulating cCREs detected in at least two contexts. Strong transcriptional activity (|log_2_(RNA/DNA)|>1|) was detected in 2-5% of active cCREs per condition, predominantly corresponding to upregulating cCREs (94% on average). Overall, 283 cCREs (~7%) showed strong upregulatory activity and 36 (<1%) showed strong downregulatory activity in at least one condition (Extended Data Fig. 3a).

**Fig. 2.**
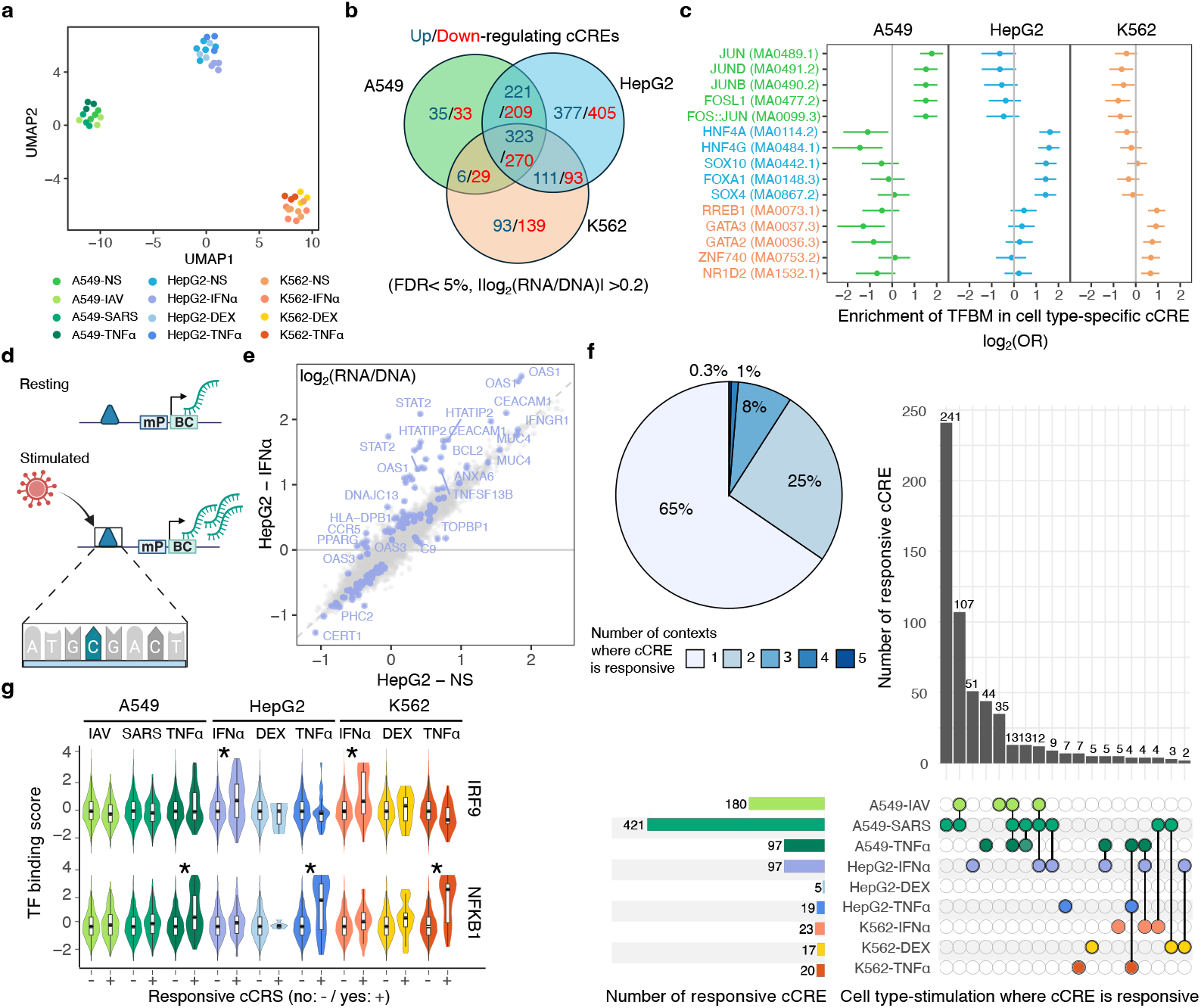
lentiMPRA reveals context-specific activity of introgressed cCREs. **a**, UMAP of transcriptional profiles across all replicates, colored by cell type and condition. **b**, Overlap of active cCREs across cell types (FDR<5%, unstimulated condition). Numbers are shown separately for upregulating (blue) and downregulating (red) cCREs. **c**, Enrichment (log_2_ odds ratio) of cell type-specific enhanced cCREs in transcription factor binding motifs (TFBM). For each cell type, only the five TFs with the strongest enrichment (based on lower bound of 95 % confidence interval) are shown. **d**, Schematic of a responsive cCRS exhibiting increased activity upon stimulation. **e**, Comparison of cCRS activity between IFN-α-stimulated and resting hepatocytes. For highly responsive cCRS, the nearest immune-related gene is reported. **f**, Sharing of responsive cCREs (FDR<5%) across cell types and stimuli. Top left: pie chart showing the proportion of context-specific versus shared responsive cCREs. Bottom left: distribution of responsive cCREs across contexts (i.e. cell types × stimuli). Right: UpSet plot showing the frequency of responsive cCREs across combinations of contexts (only combinations with ≥2 cCREs are shown). **g**, Enrichment of responsive cCREs in sequences with high binding affinity. *BH-adjusted Wilcoxon rank-sum tests (*P*<0.001).

To investigate the regulatory architecture underling these effects, we examined chromatin states, transcription factor binding motifs (TFBMs), and genomic localization. Relative to inactive elements, strongly upregulating cCREs were enriched (as defined by ChromHMM^54^) at active transcription starting sites (TSS) and TSS-flanking regions (OR>4.4, Fisher’s exact adjusted *P*<1.9×10^-4^), other upregulating cCREs at enhancers and bivalent or poised TSSs (OR>1.5, adjusted *P*<0.009), and downregulating cCREs at bivalent enhancers and Polycomb-repressed regions (OR>2.0, adjusted *P*<0.009) (Extended Data Fig. 3b and Supplementary Table 1). Distinct classes of cCREs also displayed divergent TFBM architectures (Extended Data Fig. 3c,d). Upregulating cCREs were enriched for ETS family motifs (e.g., ETV1 or EHF; OR>2.3, adjusted *P*<1.1×10^-7^) and AP-1-related motifs (e.g., FOS::JUN; OR=2.0, adjusted *P*<5.5×10^-5^), whereas downregulating cCREs preferentially harbored HOX-related motifs (e.g., HOXB4 and HOXB2; OR>1.5, adjusted *P*<0.0007) and binding sites for transcriptional repressors (e.g., SNAI1 and TBX2; OR>1.4, adjusted *P*<0.01). These elements further differed in genomic positioning relative to active TSS (CAGE-data^55^): upregulating cCREs predominated within 2kb, whereas downregulating cCREs were enriched 2-50 kb away (Extended Data Fig. 3e). Together, these results indicate that our lentiMPRA captures direction-specific regulatory activity associated with specific chromatin environments and local regulatory contexts.

### Context-specific activity of introgressed cCREs

To identify cell type-specific regulatory activity, we tested each cCRE for differential activity between cell types while adjusting for plasmid library and lentivirus batch (Methods). cCREs were classified as cell type-specific when activity was higher in one cell type relative to the other two (FDR<5%, |log_2_FC|>0.2), identifying 311, 365, and 1,009 cell type-specific cCREs in A549, HepG2, and K562 cells, respectively (Supplementary Table 2). Although regulatory activity was often shared (55%; Fig. 2b), strong cCREs showed greater specificity (cell type-specific cCRE OR=2.9, Fisher’s exact *P*<1.1×10^-12^), with only 33% active in ≥2 cell types (Supplementary Fig. 4). Consistent with lineage-specific regulatory programs, TFBM analysis revealed enrichment for hepatocyte nuclear factors such as HNF4A and HNF4G in HepG2-specific cCREs, and hematopoietic regulators such as GATA2 and GATA3 in K562-specific cCREs (Fig. 2c and Supplementary Table 2).

We next examined how stimulation affects regulatory activity (Fig. 2d,e). Across conditions, 605 cCREs showed stimulation-responsive activity (FDR<5%, |log_2_FC|>0.2; Supplementary Fig. 5 and Supplementary Table 2). IFN-α, TNF-α, and DEX predominantly increased cCRE activity (53-83% of responsive elements), whereas viral stimulation with SARS-CoV-2 or IAV led almost exclusively to decreased activity (>98%), consistent with global transcriptional shutdown in infected cells (Extended Data Fig. 4 and Supplementary Note 4). Excluding viral conditions, 21% of responsive cCREs were inactive at baseline, while the remainder were predominantly upregulating cCREs (49%), with a smaller fraction of downregulating cCREs (29%). Notably, most responsive cCREs (65%) were restricted to a single cell type and stimulus combination (Fig. 2f). Even under relaxed replication thresholds (empirical *P* [*P*_*emp*_]<0.05), responsiveness remained highly context-dependent, with only 16% of cCREs shared across ≥5 conditions (Extended Data Fig. 5). Given that responsive cCREs are expected to be enriched for targets of key immune TFs, we compared binding of 595 TFs between responsive and non-responsive cCREs (810 JASPAR core TFBMs^56^; Supplementary Table 2). IFN-α- and TNF-α-responsive cCREs were enriched for IRF9 and NF-κB targets, respectively, across cell types (Wilcoxon rank-sum *P*<1.4 × 10^-5^, Fig. 2g), supporting bona fide immune activation rather than technical variability. By contrast, DEX-responsive cCREs showed no enrichment for glucocorticoid receptor (NR3C1) motifs (*P*>0.36), suggesting predominantly indirect regulatory effects. These results demonstrate that the regulatory impact of introgressed loci is highly dependent on cellular and immune context.

### lentiMPRA identifies functional introgressed alleles across diverse populations

Having defined the activity landscape of cCREs, we next assessed how individual introgressed alleles modulate regulatory output (Fig. 3a). Allele-specific effects were quantified using the MPRAnalyze framework^57^ by comparing the activity of introgressed and non-introgressed alleles at each SNV across cell types and stimulation conditions (Supplementary Fig. 6). Across active cCREs, we identified 689 expression-modulating variants (emVars) showing allele-specific effects in at least one context (FDR<5%,|log_*2*_FC|>0.2, Supplementary Table 3), with strongly upregulating cCREs harboring the highest fraction (Supplementary Fig. 7). Introgressed alleles were equally likely to increase or decrease cCRE activity, and effect directions were concordant with eQTL^58^ and baseline MPRA^49^datasets (Spearman’s ρ=0.5, Supplementary Note 5). Although emVars spanned a continuum of regulatory effects, 90 variants showed strong modulation (|log_2_FC|>0.5) across cell types (Fig. 3b and Extended Data Fig. 6a).

**Fig. 3.**
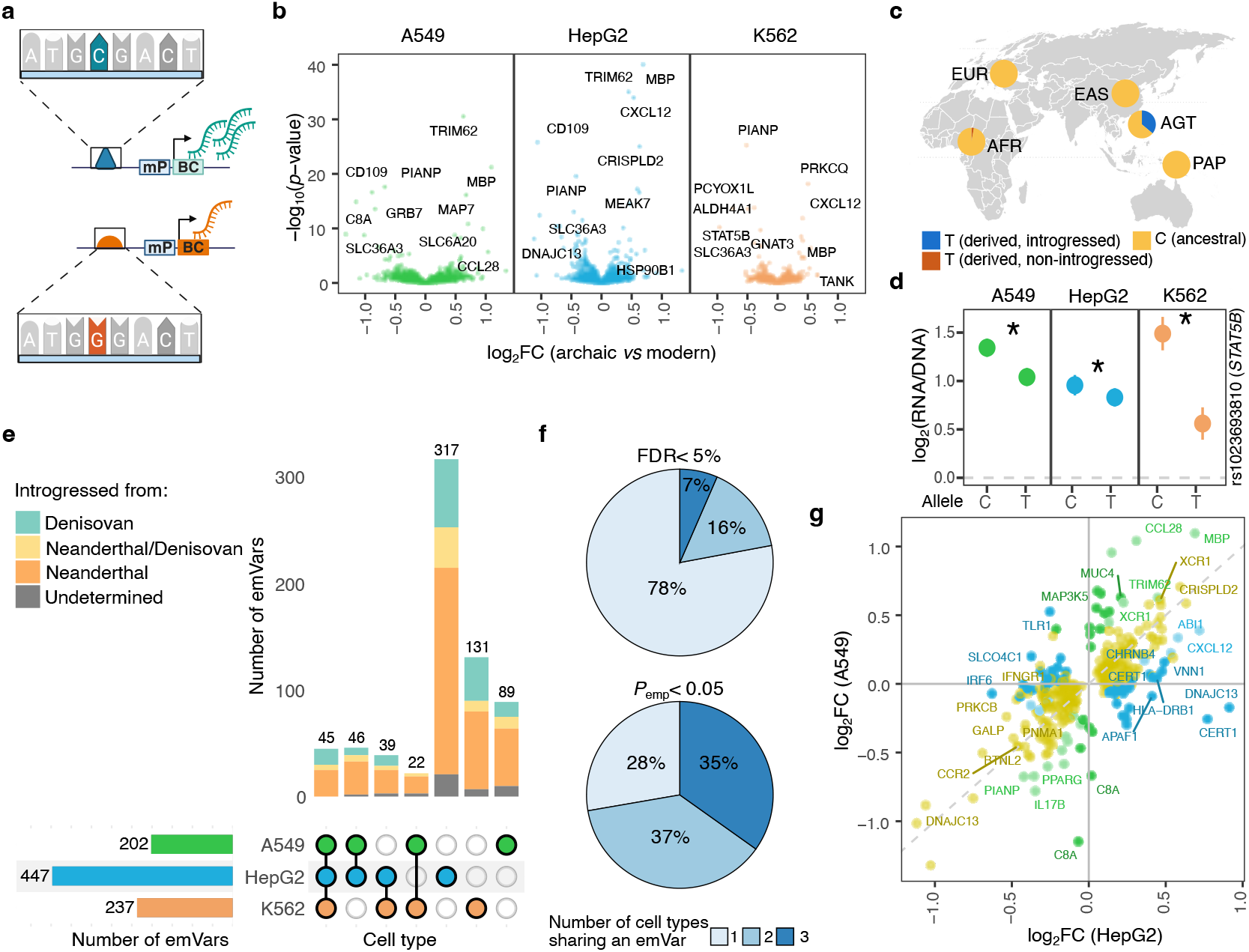
Introgressed alleles exhibit cell type-specific effects on gene expression. **a**, Schematic of an emVar in which the C allele is associated with increased gene expression. **b**, Volcano plots showing log_2_ fold change (log_2_FC) between alleles and –log_10_(*P* value) for differential expression across all three cell types (unstimulated condition). For emVars with the strongest *p*-value or largest effect sizes, the gene with the nearest TSS is indicated. **c**, Worldwide frequencies of rs1023693810 alleles in the SGDP panel. Introgressed haplotypes (absent in Africa and carrying the linked archaic-derived rs113642041-G allele) are shown in blue and distinguished from non-introgressed haplotypes (carrying rs1023693810-T and rs113642041-A alleles) shown in red. **d**, Estimated cCRE activity of each allele of rs1023693810 across the three tested cell types (basal condition only). Error bars indicate 95% confidence intervals. **e**, UpSet plot showing sharing of emVars across the three cell types. Colors reflect the inferred archaic source of the introgressed haplotype. **f**, Pie charts showing the proportion of emVars that are context-specific or shared across cell types at the detection level (top, FDR<5%) and replication level (bottom, *P*<0.05). **g**, Comparison of emVar effect sizes between A549 and HepG2 cells. Only emVars reaching FDR<0.05 in at least one cell type under unstimulated conditions are shown. Colors indicate sharing patterns: shared (yellow, *P*_emp_<0.05 in both cell types, *P*_GxCell-type_>0.05), light blue/green: HepG2/A549-enhanced, *P*_emp_<0.05 in both cell types, *P*_GxCell-type_<0.05, same direction, dark blue/green: HepG2/A549-specific (FDR<0.05 in one cell type and *P*_emp_>0.05 in the other).

A substantial fraction of strong-effect emVars (80%) occurred in populations underrepresented in genomic studies, including Papuans, Agta hunter-gatherers from the Philippines, and South Asians (OR=1.76, Fischer’s exact *P*=0.04). Some emVars reduced regulatory activity at immunity-related loci. For example, rs1023693810-T (36% in Agta) at *STAT5B* decreased the activity of a distal enhancer (EH38E3223888, SCREEN database^59^) across cell types, most prominently in K562 cells (log_2_FC=-0.50; Fig. 3c,d). According to Encode Hi-C data^59^, this enhancer physically interacts with up to ten genes, including *STAT5B, STAT5A*, and *STAT3*. Other emVars increased regulatory activity, such as the ancestral allele rs57870697-G, reintroduced into Papuans by Neanderthals (56%), at a *STAT2*-proximal enhancer (EH38E3020497) in HepG2 cells (Extended Data Fig. 7). These examples illustrate how MPRA can resolve the regulatory effects of population-enriched introgressed alleles, uncovering functional variation that remains largely inaccessible to standard genomics studies.

### Introgressed alleles exhibit cell type- and stimulus-specific regulatory effects

We first quantified the extent to which emVars were shared across cellular contexts. Although most emVars (~78%) were initially detected in a single cell type, they significantly overlapped across cell types (OR>4.9, *P*<1.5×10^-23^), (Fig. 3e). Under a more relaxed threshold (*P*_*emp*_<0.05), 72% replicated in at least another cell type, and effect sizes were strongly correlated across conditions (Spearman’s ρ=0.40-0.86, *P*<2.2×10^-16^), indicating that limited power inflates apparent specificity (Fig. 3f, Extended Data Fig. 6b, Supplementary Fig. 8 and Supplementary Note 6). Nevertheless, interaction tests showed that 41% of emVars had significantly different effect sizes between cell types (MPRAnalyze likelihood ratio test, permutation-based FDR<5%, Fig. 3g and Supplementary Table 3), indicating that quantitative differences in regulatory effect also contribute to context specificity.

We then examined how immune and infectious stimuli reshape the regulatory effects of introgressed alleles (Fig. 4a). Among the 689 identified emVars, 180 were detected exclusively after stimulation (*P*_emp_>0.05 at baseline), whereas only 29 were specific to unstimulated cells (*P*_emp_>0.05 across stimulated conditions; Supplementary Table 4).Genotype-by-stimulation interaction tests further identified 94 emVars whose effects were revealed or modified by stimulation (FDR_GxE_<5%, |Δlog_2_FC|>0.2; Supplementary Table 4). These effects were highly context-dependent: 85% of responsive emVars were observed in a single cell type and stimulation condition, and even under a marginal interaction threshold (*P*_emp_ <0.05), 43% retained this dual specificity (Fig. 4b,c and Extended Data Fig. 8). The strongest responsive emVar was rs57784925-G at *TRIM62*, an ancestral allele reintroduced into most non-Africans by Neanderthals (24-31% in Eurasia) but rare in the Pacific (Fig. 4d). At baseline, the introgressed G allele specifically repressed transcription in K562 cells, but upon TNF-α stimulation, it markedly increased responsiveness particularly in K562 cells (Δ log_2_FC=0.31-0.76; Fig. 4e). Sequence analysis indicated that the G allele strengthens an NF-κB binding motif at this locus (Fig. 4f and Supplementary Table 4). Consistent with an immunological role, phenome-wide association analysis linked rs57784925-G to improved pulmonary function and reduced asthma risk (Student’s *P*<6 ×10^-4^). Together, these results show that immune stimulation frequently reveals regulatory effects of introgressed alleles, highlighting environmentally responsive regulatory variation as an important component of the functional legacy of archaic introgression.

**Fig. 4.**
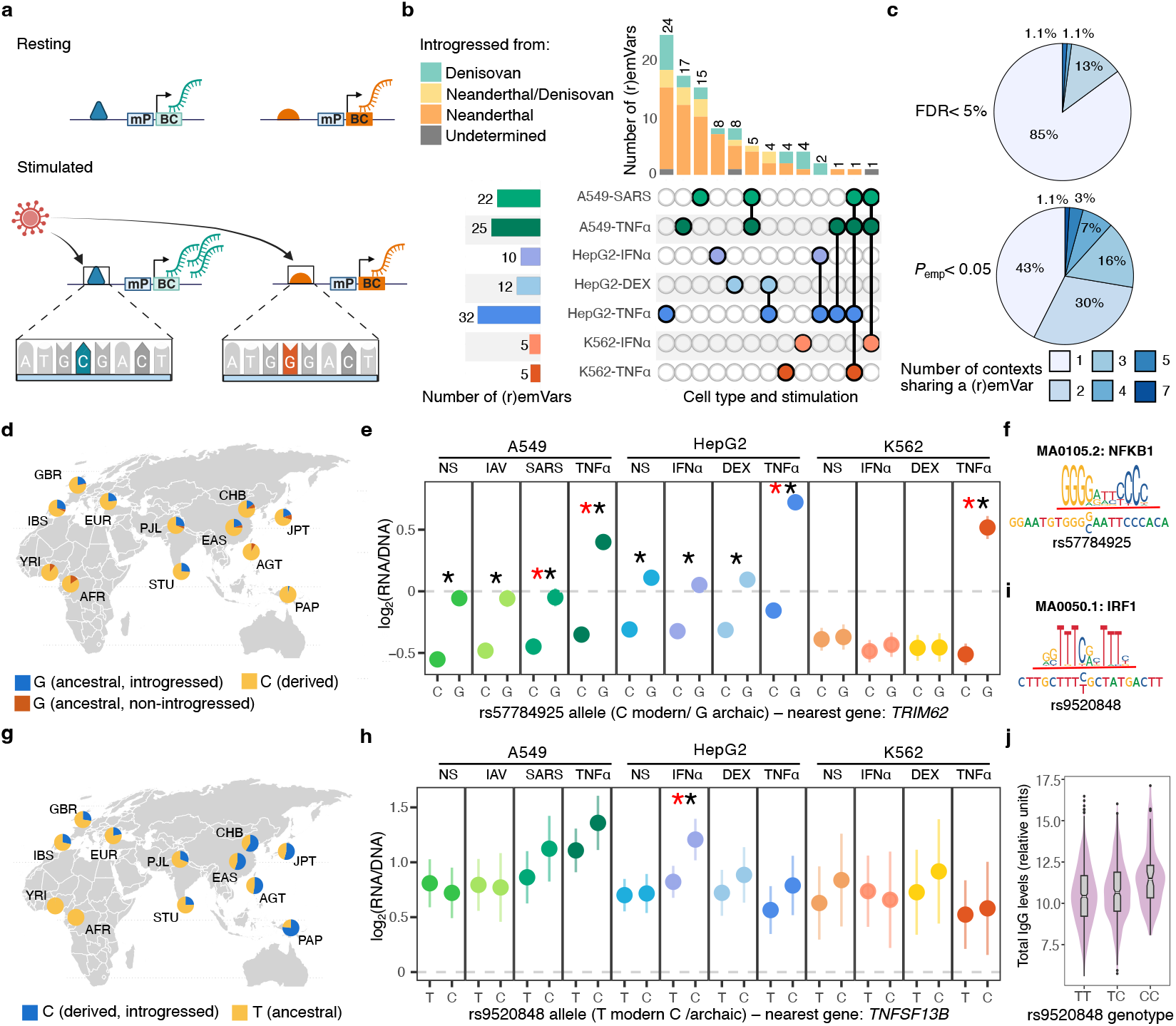
Adaptively introgressed alleles exert stimulation-dependent effects on gene expression. **a**, Schematic of a responsive emVar, (r)emVar, in which the C allele increases gene expression specifically upon viral stimulation. **b**, Sharing of responsive emVars (FDR<5%) across cell types and stimuli. UpSet plots show their distribution across contexts (cell type×stimuli) and combinations thereof. **c**, Pie charts showing the proportion of responsive emVars that are context-specific or shared across conditions at the detection level (top; FDR<5%) and replication level (bottom; *P*<0.05). **d**,**g**, Worldwide allele frequencies of rs57784925 (**d**) and rs9520848 (**g**) in the SGDP and 1KGP panels. For rs57784925-G, introgressed haplotypes (absent in Africa and carrying the linked archaic-derived rs12035312-G allele) are shown in blue, and distinguished from non-introgressed haplotypes (carrying rs57784925-G and rs12035312-A allele) shown in red. **e, h**, Estimated cCRE activity of each allele of rs57784925 (**e**) and rs9520848 (**h**) across all tested conditions. Error bars indicate 95% confidence intervals. Red stars denote responsive emVars (FDR<5%); black stars denote emVars (FDR<5%). **f, i**, Predicted impact of rs57784925 (**f**) and rs9520848 (**i**) on TFBMs for NF-κB and IRF1. **j**, Association of the rs9520848-C allele with increased total IgG levels in the *Milieu Intérieur* cohort.

### Context-dependent effects of introgressed alleles contribute to disease risk

To assess the disease relevance of emVars, we systematically intersected them with GWAS signals. On average, 33% of emVars were associated with at least one trait (Supplementary Table 4). Notably, 12% of trait-associated emVars lacked reported links to gene expression or splicing in the eQTL Catalogue^58^, highlighting the added resolution of lentiMPRA for resolving regulatory mechanisms underlying GWAS loci. Among the strongest associations were two Denisovan-introgressed alleles at *IKZF3* (rs1510475-C and rs34016964-T, 53% in Papuans; Extended Data Fig. 9a,b), prioritized for their long-range interactions with the *ORMDL3* promoter in T cells. These variants, which lie on the same archaic haplotype in Papuans (r^2^ >0.8) and are also found in Europe on a non-archaic haplotype segregating at low frequency (3%), are associated with altered *IKZF3* expression in T, NK, and B cells, and increased risk of systemic lupus erythematosus^60^. Yet, their regulatory effects diverge: rs1510475-C decreased activity across all cell types, whereas rs34016964-T increased reporter expression specifically in K562 cells (Extended Data Fig. 9c,d).

Intersecting the 94 responsive emVars with immune- and infection-related GWAS loci identified 17 overlaps (Supplementary Table 4). One notable example, rs9520848-C, displayed a pronounced IFN-α-dependent effect in HepG2 cells (Fig. 4g-j). The introgressed C allele enhances IRF1 binding to a distal enhancer (EH38E1696591) located 34 kb downstream of *TNFSF13B*, encoding the B-cell activating factor BAFF (Fig. 4i). This allele lies on a 21.9-kb haplotype shared across archaic genomes segregating at high frequency in modern non-Africans (22-77%; Fig. 4g), consistent with strong positive selection. In line with its regulatory effects, rs9520848-C is associated with increased monocyte counts (UK biobank, GCST90692929)^61^ and elevated *TNFSF13B* expression in MAIT cells^29^. Phenome-wide association analysis further linked this allele to increased plasma IgG levels (Fig. 4j) and revealed nominal associations with systemic lupus erythematosus (Extended Data Fig. 9e and Supplementary Table 4). These results show that context-dependent regulatory effects of introgressed alleles contribute to variation in immune traits and disease susceptibility, providing a functional link between archaic introgression and disease risk.

### Coordinated regulatory effects reveal adaptive expression phenotypes

To test whether Neanderthal and Denisovan alleles differ in regulatory potential, we estimated, for each condition, the proportion of variants with non-zero effects on gene expression (*π*_1_) among introgressed alleles from each archaic lineage (Fig. 5a and Supplementary Table 5). Overall, Neanderthal- and Denisovan-introgressed variants were similarly likely to influence gene expression, except in A549 cells, where Neanderthal variants more frequently altered expression following IAV and TNF-α stimulation (OR=1.7 and 1.4; BH-adjusted Fisher’s exact *P*=0.003 and 0.045). Stratification by effect direction further showed that Denisovan alleles in Papuans were less likely to decrease expression under these conditions than introgressed alleles worldwide, suggesting purifying selection against expression-decreasing variants in this population (Fig. 5b and Supplementary Fig. 9).

**Fig. 5.**
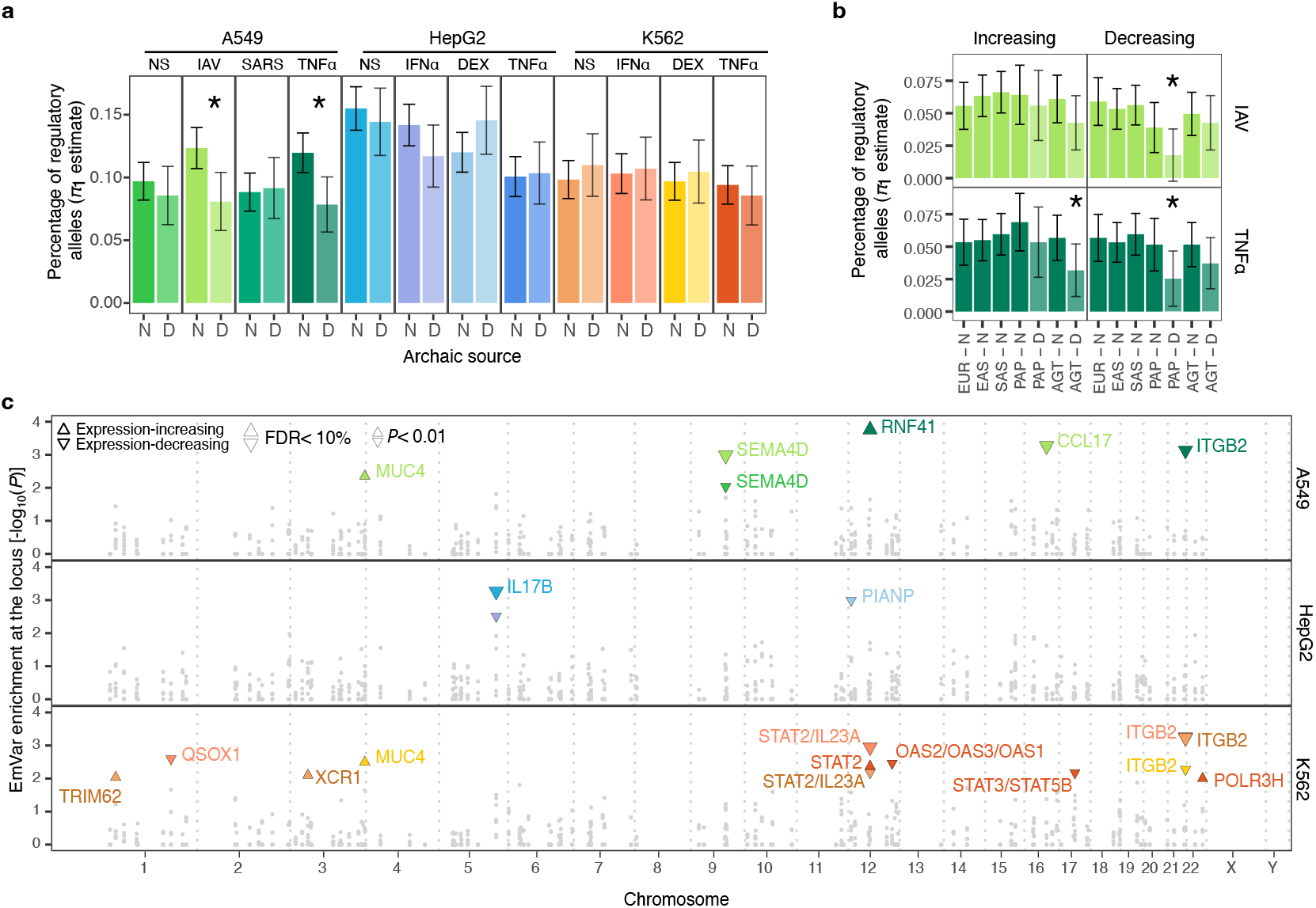
Coordinated regulatory effects of adaptively introgressed alleles. **a**, Estimated proportion (π_1_) of Neanderthal (N) and Denisovan (D) alleles that affect expression across all tested conditions. Bars indicate 95% bootstrap confidence intervals. Colors denote cell type and condition. *BH-adjusted Fisher’s exact test *P*<0.05. **b**, Breakdown of proportion of regulatory alleles by population, archaic source, and direction of effect. *Fisher exact test *P*<0.05. **c**, Local enrichment of archaic haplotypes in regulatory variants. Upper and lower triangles indicate enrichment of expression-increasing or expression-decreasing alleles, respectively (Fisher’s exact *P*<0.01). Colors denote the condition in which the enrichment is observed. Loci passing 10% FDR are indicated by larger triangles. For each significant enriched haplotype, nearby immune-relevant genes (within 10kb) showing suggestive enrichment (Fisher’s exact *P*<0.05) are reported.

We next sought to identify adaptive expression phenotypes at introgressed immune loci. Under the poison-antidote model, adaptive introgression is expected at loci that protected archaic groups from long-standing endemic pathogens, which were subsequently introduced into modern humans^20^. At such loci, we expect repeated fixation of alleles with coordinated regulatory effects in archaic hominins, generating local regulatory divergence between introgressed and non-introgressed haplotypes. To test this prediction, we assessed whether introgressed haplotypes are locally enriched for expression-increasing (*P*_emp_<0.05, log_2_FC>0) or expression-decreasing (*P*_emp_<0.05, log_2_FC<0) alleles relative to genome-wide expectations. We identified 13 archaic haplotypes with marginal enrichment (Fisher’s exact *P*<0.01), five of which remained significant at 10% FDR threshold after BH correction (Fig. 5c and Supplementary Table 5). The strongest signal corresponded to a Neanderthal haplotype at the *STAT2* locus^24^, which was enriched for alleles increasing expression in TNF-α-stimulated A549 cells (OR=7.1, Fisher’s exact *P*=0.0002) and decreasing expression in IFN-α-stimulated K562 cells (OR=5.9, Fisher’s exact *P*=0.001).

To refine candidate gene targets, we repeated the analysis within 10-kb windows surrounding genes located within these haplotypes (Supplementary Table 5). *STAT2* and *IL23A* showed the strongest enrichment of expression-decreasing alleles in IFN-α-stimulated K562 cells (OR>10.3, Fisher’s exact *P*<0.014), whereas *RNF41*, encoding the antiviral ubiquitin ligase NRDP1 (ref.^62^), showed the strongest enrichment of expression-increasing alleles in TNF-α-stimulated A549 cells (OR=10.1, Fisher’s exact *P*=0.025). Additional enrichments of expression-decreasing alleles were detected near *CCL17* (OR=115, *P*=6.6×10^-5^) and *SEMA4D* (OR=7.5, *P*=0.001) in IAV-stimulated A549 cells, as well as near *IL17B* in HepG2 (OR=5.4, *P*=0.005) and *ITGB2* in A549 and K562 cells (OR>11.4, *P*<0.004).Together, these results identify loci where introgressed alleles exert coordinated, condition-specific effects on gene expression, highlighting putative adaptive expression phenotypes shared between archaic and modern humans.

### Neanderthal COVID-19 risk haplotype drives TNF-α responsiveness in lung epithelium

Finally, we asked whether our data could clarify the cellular and molecular effects of Neanderthal haplotypes associated with COVID-19 risk^13,27^. A Neanderthal haplotype on chromosome 12 encompassing the *OAS1-3* cluster reaches high frequencies across mainland Eurasia (15-36%) and has been associated with reduced COVID-19 severity^27^. Our lentiMPRA included 91 putatively introgressed alleles at this locus, 59 of which overlapped the 95% credible set for COVID-19 risk (Extended Data Fig. 10a,b). We identified 20 emVars, including 12 associated to COVID-19 phenotypes (Extended Data Fig. 10c and Supplementary Table 6). The strongest regulatory effect at this locus was observed for rs1859333-C>T in IAV-stimulated A549 cells. This variant disrupts a cCRE with strong downregulating activity (EH38E3042014) located 5kb upstream of *OAS3* and predicted to interact with *OAS1* and *OAS3* in lung tissue (Extended Data Fig. 10d). By contrast, the only allele with a significant effect in SARS-CoV-2-stimulated A549 cells was rs4766674-G in *OAS1*, which increased transcriptional output across all cell types and conditions despite lacking an associated ENCODE cCRE annotation (Extended Data Fig. 10e).

We next examined the Neanderthal-introgressed chromosome 3 locus encompassing multiple genes (*LIMD1, SACM1L, SLC6A20, LZTFL1, CCR9, FYCO1, CXCR6, XCR1, CCR1* and *CCR3*), which has been associated with increased risk of COVID-19 hospitalization^13^. The core 49-kb haplotype overlapping the 95% credible set for disease severity displays marked population differences, being nearly absent in East Asians (<1%), present in Europeans (4-7%), and common in South Asians (25-30%), Papuans (33%) and the Agta (53%) (Fig. 6a,b). Our lentiMPRA contained 230 Neanderthal-introgressed variants across this locus, among which we identified 49 emVars (Supplementary Table 6). Notably, among the four emVars overlapping the core haplotype, rs17713054-A showed the strongest regulatory effect among all 3,838 variants tested in the lentiMPRA (|log_2_FC|=1.69, FDR<0.001), with pronounced cell type specificity. Located within a strong enhancer (EH38E2197611), this allele increased transcription by ~2-fold specifically in unstimulated A549 cells (|log_2_FC|>0.93, FDR<0.002), and its effect was further amplified following TNF-α stimulation (>3-fold increase, |Δ log_2_FC|=0.76, FDR_G×E_=0.004; Fig. 6c). Given the association between circulating TNF-α levels and COVID-19 severity^63^, these results implicate rs17713054-A as a key functional driver of the chromosome 3 risk haplotype through enhanced inflammatory responsiveness in lung epithelial cells.

**Fig. 6.**
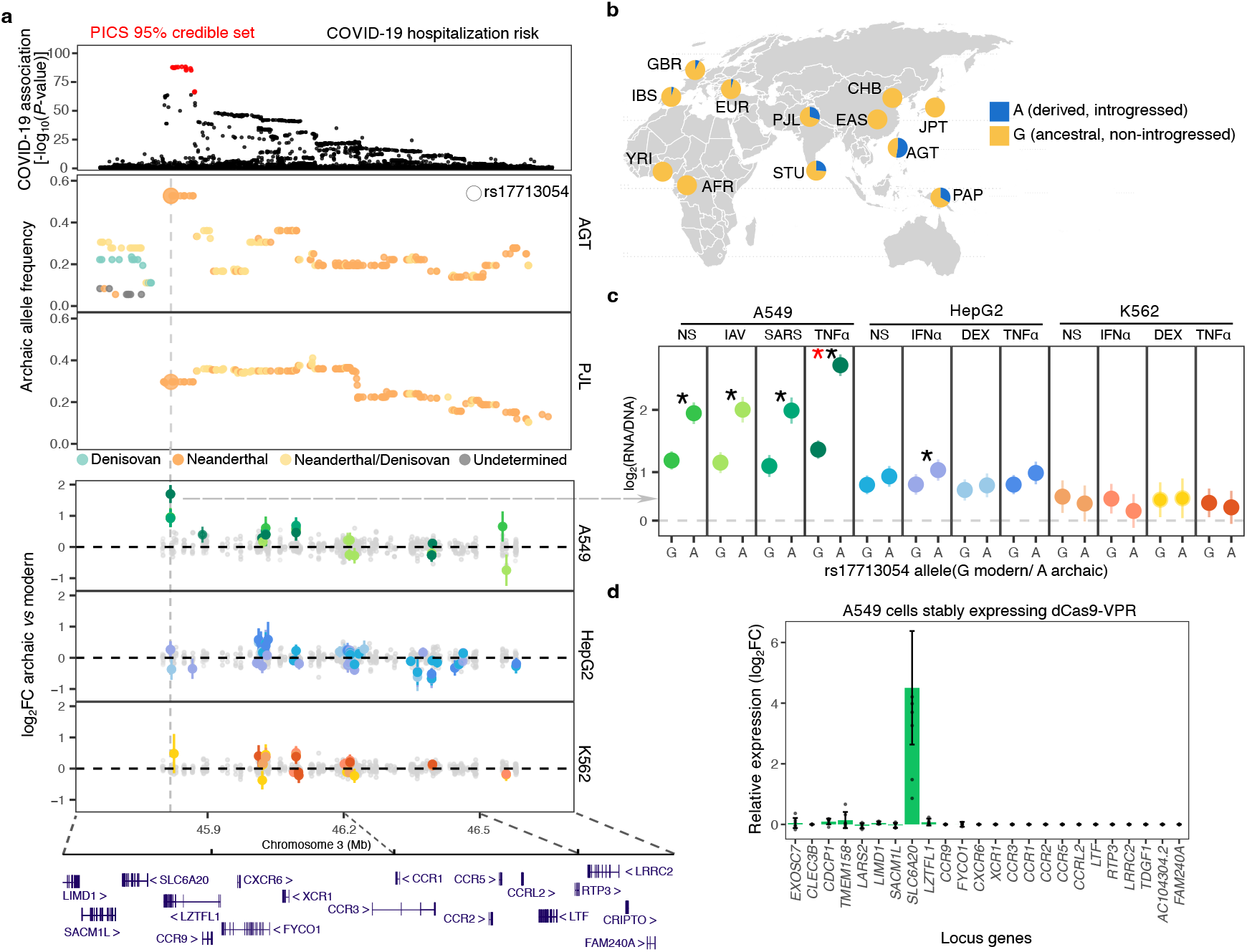
Introgressed alleles at the COVID-19 risk locus confer TNF-α-responsiveness in lung epithelial cells. **a**, Overview of introgressed alleles at the chromosome 3 COVID-19 risk locus and their regulatory effects. Top: locusZoom plot of association with COVID-19 hospitalization risk (GCST90134600), variants in the 95% credible set are highlighted in red. Upper middle: frequency, genomic localization, and archaic source of introgressed alleles in Agta (AGT) and Punjabi from Lahore (PJL). Alleles are colored by archaic match: Neanderthal (orange), Denisovan (teal), both (yellow) or neither (grey). Lower middle, MPRA-inferred regulatory effects. For each variant, the log_2_FC with 95% confidence intervals is reported across all cell types and stimuli; non-significant effects are shown in grey, significant emVars (FDR<5%, |log_2_FC|>0.2) are colored by condition as in (**c**). Bottom, protein-coding gene annotation at the locus. **b**, Worldwide allele frequencies of rs17713054 in 1KGP and SGDP panels, with archaic genotypes shown for reference. **c**, Estimated cCRE activity of each allele of rs11713054 across all tested conditions. Error bars indicate 95% confidence intervals. Red stars denote responsive emVars (FDR<5%); black stars denote emVars (FDR<5%). **d**, Fold change in expression of protein-coding genes at the chromosome 3 locus following CRISPR activation of the rs17713054-overlapping enhancer. Bars indicate 95% confidence intervals; points represent individual replicates.

To identify the target gene(s) of the rs17713054 enhancer, we performed CRISPR activation in A549 cells expressing dCas9-VPR^64^, using a pool of three sgRNA targeting the enhancer (Methods). qPCR revealed activation of *SLC6A20*, but not the neighboring genes *LZTFL1* and *CXCR6* (Supplementary Fig. 10), despite previous in silico studies proposing the latter genes as candidate targets^65,66^. Consistently, bulk RNA-seq comparing cells with rs17713054-targeting sgRNAs versus control sgRNA showed strong and highly specific upregulation of *SLC6A20* (log_2_FC=4.5, FDR<0.002; Fig. 6d), with no detectable effects on adjacent genes. Together, these results functionally resolve the chromosome 3 Neanderthal risk haplotype by demonstrating that an enhancer overlapping rs17713054 directly regulates *SLC6A20* in lung epithelial cells, thereby identifying the molecular effector underlying the strongest genetic risk locus for severe COVID-19.

## Discussion

In this study, we assessed the functional impact of common immunity-related archaic alleles across basal and stimulated conditions in three cell lines. Our lentiMPRA framework enabled the detection of active cis-regulatory elements with direction-specific effects across a wide variety of contexts. Within these elements, we identified 689 variants for which introgressed and non-introgressed alleles displayed different regulatory activity. Importantly, the effects of 94 of these emVars were either revealed or significantly altered upon immune stimulation or viral exposure, underscoring the importance of assaying regulatory variation in biologically relevant contexts to understand its phenotypic consequences.

Expanding functional analyses beyond European-ancestry populations enabled us to assign function to numerous population-specific introgressed alleles. A notable example is a high-frequency Denisovan haplotype on chromosome 17 in the Agta population that harbors multiple expression-decreasing alleles, including rs1023693810-T at *STAT5B*. This variant modulates the activity of a constitutive enhancer linked to a gene central to growth hormone signaling and multiple cytokine pathways^67^. Homozygous loss-of-function mutations in *STAT5B* cause severe growth retardation, immune dysregulation, and anemia^67^, whereas heterozygous variants reduce height^68^. Denisovan-mediated attenuation of *STAT5B* enhancer activity may therefore contribute to the shorter stature of the Agta, a phenotype proposed to be adaptive in tropical rainforest hunter-gatherer environments^69^. More broadly, these findings illustrate how studying underrepresented populations can uncover regulatory variants—here introduced through archaic admixture—contributing to population-specific phenotypes.

Numerous emVars also act in a cell type- or stimulus-specific manner, with one-third overlapping eQTLs studies and GWAS loci. Notably, the strongest responsive emVar is Neanderthal-introgressed, occurs at high frequency (~28%) in mainland Eurasians, and maps to the *TRIM62* locus, where its effect is amplified after TNF-α stimulation across all cell types. *TRIM62* encodes an E3 ubiquitin ligase involved in NF-κB, AP-1, and interferon pathways^70^, and Trim62^-^/^-^ mice display impaired pathogen clearance, increased mortality to fungal infection, and heightened susceptibility to colitis^71^. These observations suggest that the introgressed rs57784925-G allele may confer protection against infection by enhancing expression in response to TNF-α, a cytokine released during acute inflammatory responses. Similarly, the archaic-derived rs9520848-C variant (22-77%) increases activity of a BAFF-interacting enhancer in an IFN-α-dependent manner in liver cells. Given the central role of BAFF in promoting B-cell survival and antibody production^72,73^, and the efficacy of BAFF-antagonist therapies in systemic lupus erythematosus^74^, this variant may reflect an evolutionary tradeoff between pathogen defense and autoimmune risk. Collectively, these findings demonstrate that profiling regulatory activity under immune stimulation reveals clinically relevant molecular phenotypes that would otherwise go undetected.

Our study also clarifies the mechanism linking the chromosome 3 Neanderthal haplotype to COVID-19 severity^13^. We identify rs17701354-G>A, the lead variant in COVID-19 meta-analyses^75^, as a highly context-specific regulatory allele that strongly increases enhancer activity in lung epithelial cells, particularly following TNF-α stimulation. This effect was not detected in a previous K562-based MPRA^44^, underscoring the importance of cellular context for resolving disease-associated regulatory mechanisms. CRISPR activation experiments identified *SLC6A20* as the target gene of the rs17701354-overlapping enhancer, with the Neanderthal-derived risk allele specifically increasing *SLC6A20* expression, consistent with evidence that loss of *SLC6A20* improves A549 cell survival during SARS-CoV-2 infection^66^. *SLC6A20* encodes the sodium/imino-acid transporter 1 (SIT1), which interacts with the SARS-CoV-2 receptor ACE2^76^. Interestingly, SIT1 overexpression retains ACE2 in the cytosol, reducing its cell-surface levels^77^. These observations raise the possibility that rs17701354-A increases COVID-19 severity not by enhancing viral entry, but through ACE2 sequestration and disruption of the renin-angiotensin system, thereby exacerbating cytokine storm and acute respiratory distress^78,79^. Together, these findings support *SLC6A20* as the primary effector gene underlying the Neanderthal-derived COVID-19 risk haplotype, refining previous models that implicated neighboring genes including *LZTFL1, CXCR6, CCR1*, and *CCR5* (ref.^44,65,66^).

Beyond rs17701354, most emVars within this Neanderthal haplotype lie outside the 95% credible set for COVID-19 severity and likely affect additional immune phenotypes, including decreased *CCR5* expression and reduced HIV susceptibility^80^. We further observed enrichment of expression-increasing alleles in resting and IFN-α-stimulated K562 cells (OR>2.1, Fisher’s exact *P*<0.007; Supplementary Table 5), particularly near *XCR1* (OR>5.7, Fisher’s exact *P*<0.008), a key receptor in dendritic cell antigen cross-presentation and cytotoxic immune responses^81^. These patterns suggest that selection on beneficial immune functions contributed to the maintenance of the haplotype in present-day populations, with the COVID-19 risk allele increasing in frequency through genetic hitchhiking.

Our study has several limitations. First, focusing on high-frequency immunity-related variants limits inference about the broader regulatory potential of neutral or negatively selected archaic alleles. Second, bulk lentiMPRA cannot resolve variants active in rare or transient cellular states, including infected cells with varying viral loads, highlighting the potential of single-cell MPRAs^82^ to dissect infection-specific regulatory programs. Finally, despite the diversity of cellular contexts examined here, additional cell types, environmental perturbations, and systematic target-gene mapping will be required to fully characterize the phenotypic consequences of archaic introgression beyond immunity.

Despite these limitations, our study provides a functional framework for understanding how Neanderthal- and Denisovan-inherited variants shape transcriptional responses to immune challenges in present-day humans. More broadly, it illustrates how context-aware functional genomics can bridge the gap between statistical associations and molecular mechanisms, enabling systematic dissection the evolutionary and biomedical consequences of archaic introgression.

## Methods

### Generation of archaic-introgressed variant dataset

#### Whole-genome sequence datasets

To systematically detect archaic-introgressed haplotypes across diverse human populations, we applied a combination of two complementary methods: conditional random fields (CRF)^83,84^ and the S’ statistic^85^) to whole-genome sequencing data from (i) the 1000 Genomes Project (1KGP Phase 3, ftp://ftp.1000genomes.ebi.ac.uk/vol1/ftp/release/20130502/)^86^ and (ii) a previously compiled high coverage (30×) dataset integrating sequences from the SGDP and populations from Near Oceania^87^, hereafter referred to as to *SGDP-Pacific* dataset. For the 1KGP dataset, we selected two populations from each major continental group, excluding South America due to high admixture that could bias LD-based detection of archaic introgression. Specifically, we used British from England and Scotland (GBR) and Iberians from Spain (IBS) for Europe; Han Chinese from Beijing (CHB) and Japanese from Tokyo (JPT) for East Asia; Punjabi from Lahore (PJL) and Sri Lankan Tamils from the UK (STU) for South Asia. Yoruba from Ibadan, Nigeria (YRI) served as the reference African population. Analyses were restricted to bi-allelic SNVs, with duplicates removed using *bcftools* v.1.9 (ref.^88^) (*view -m2 -M2 -v s* and *norm -d snp)*. For the *SGDP-Pacific* dataset, we focused on the Philippine Agta (AGT) and Papuans from New Guinea (PAP), both of which harbor substantial Denisova-related ancestry inherited through independent introgression events^87,89-91^. SGDP Africans were included as an African reference, with West Europeans and East Asians SGDP samples serving as additional continental references.

#### Archaic genomes

High-coverage VCFs for the Vindija33.19 and Altai Neanderthals, as well as the Altai Denisovan, were obtained from http://cdna.eva.mpg.de/neandertal/Vindija/VCF/. Prior to merging with modern human datasets, we systematically excluded SNVs that (i) were heterozygous, (ii) had missing genotypes, (iii) fell outside of Vindija/Altai Neanderthal and Altai Denisovan accessibility masks (downloaded from http://ftp.eva.mpg.de/neandertal/Vindija/FilterBed/), (iv) overlapped known CpG islands (obtained from UCSC Table Browser^92^), or (v) overlapped known segmental duplications (downloaded from http://hgdownload.cse.ucsc.edu/goldenPath/hg19/database/genomicSuperDups.txt.gz).

#### Detection of archaic-introgressed regions

For each dataset and population, we first applied a CRF method^83,84^ to call Neanderthal- and Denisovan-derived haplotypes. For Neanderthal introgression, the Vindija33.19 genome^93^ served as the archaic reference, with Africans (YRI or SGDP) and the Altai Denisovan genome as the non-introgressed population. For Denisovan introgression, the Altai Denisovan genome^94^ was the archaic reference, with Africans (YRI or SGDP) and Vindija33.19 as the non-introgressed population. For each target population, regions were defined as introgressed if at least one individual carried a haplotype with >90% probability of being derived from either archaic source. In parallel, we used the S’ statistic^85^ to detect introgressed segments potentially missed by the CRF, including regions derived from unknown hominins or where Neanderthal and Denisovan sequences are highly similar. Recombination rates were obtained from the 1KGP Phase 3 genetic map^86^. Regions with a S’ >150,000 were considered as introgressed, a threshold previously shown to balance sensitivity and specificity^87^.

#### Identification of high-confidence archaic-derived alleles

For each site, the ancestral state was defined as the allele present in the chimpanzee reference genome (panTro4) aligned to hg19, as downloaded from the UCSC platform^92^. The other allele was defined as derived. Sites were excluded if they were absent from the chimpanzee genome or if the chimpanzee allele differed from both the human reference and alternative alleles.

In each target population, we annotated derived alleles as putative *archaic-derived* (aSNPs) if they met the following criteria: (i) present in either heterozygous or homozygous state in the Vindija33.19 or Altai Denisova genome; (ii) absent in one of the two African reference populations (YRI or SGDP Africans) and at <2% frequency in the other; and (iii) located within an introgressed region previously identified in that population. Focusing on aSNPs with minor allele frequency >0.05, we used *plink*^95^ with the options *--r2 --maf 0*.*05 --ld-window 999999 --ld-window-kb 500 --ld-window-r2 0*.*8* to compute pairwise LD and to identify, for each aSNP, all linked aSNPs in the population of interest. The putatively introgressed haplotype was defined as the genomic interval between the most upstream and most downstream linked aSNPs. We further filtered out aSNPs where this haplotype was <10kb to reduce the risk of incomplete lineage sorting. This procedure yielded 183,652 high-confidence *archaic-derived* alleles segregating at >5% frequency in modern humans.

Beyond the direct effects of aSNPs, archaic introgression can alter human phenotypes through the reintroduction of ancient alleles that predate the split between archaic and modern humans but were lost in modern humans, either locally (in the population where introgression occurs) or globally^22,51^. To capture such “re-introgressed” alleles, we extended the set to include any allele in strong LD (r^2^>0.8) with >2 high-confidence aSNPs. This resulted in a total of 487,386 putative introgressed alleles (*common introgressed alleles*) for downstream analyses.

To define candidate adaptively introgressed haplotypes, we first extracted aSNPs detected for each archaic source and population and used the frequency distribution of their derived alleles to approximate that of the true frequency distribution of introgressed alleles. Introgressed alleles with frequencies exceeding the 95th percentile of the aSNP distribution were considered candidate targets of adaptive introgression. Estimated frequency distributions and corresponding thresholds for each population and archaic source can be found in Extended Data Fig. 1a.

#### Definition of non-overlapping introgressed regions

To define a reduced set of loci representing largely independent introgression events, introgressed alleles were grouped along the genome based on physical proximity. Specifically, we defined *introgressed regions* as sets of common introgressed alleles separated by <100 kb between consecutive variants, irrespectively of the population in which introgression was detected. This threshold corresponds to the 99.5th percentile of observed distances between consecutive introgressed alleles (median ~1kb) and is sufficient to separate independent introgressions events while grouping nearby haplotypes fragmented by recombination. Using this definition, the 487,386 common introgressed alleles were grouped in ~2300 regions, each spanning 10kb to 10.7Mb (median ~266kb) and containing a median of 85 alleles per region (range: 3-5,285).

#### Identification of archaic source at each introgressed locus

For each introgressed variant and population, the archaic source was assigned using a two-step process. First, we analyzed high-scoring S’ segments and computed, for each segment, the proportion of alleles matching Neanderthal or Denisovan genomes, as previously described^85^. Segments with a match rate >0.3 to a single hominin were assigned to that source, whereas segments with match rates >0.3 to both or <0.3 to either were classified as shared or undetermined, respectively. Second, we used the CRF-based haplotype calls^84^ to refine assignments for segments uniquely detected by this method or classified as shared/undetermined by the S’ approach. Variants were re-assigned as *Neanderthal, Denisovan*, or *shared* if the CRF could assign at least one haplotype with >90% probability to *Neanderthal, Denisova* or *both*, and kept their original assignment otherwise. For re-introgressed variants, which are not captured by the S’ method,the source was assigned from linked aSNPs (r^2^>0.8 within the same population). When assignments differed across populations, a consensus source was defined based on the population in which the allele reached the highest frequency.

#### Subset selection of immune-relevant loci

To investigate the impact of introgressed variants on immune function, we focused on a set of immune-relevant genes, defined as genes annotated under Gene Ontology term GO:0006955 (n=1,896 genes) or encoding virus-interacting proteins (VIPs) with characterized effects on viral activity (pro/antiviral)^96^ (n=119 genes, including 27 overlapping with GO:0006955). We then filtered our set of archaic-introgressed variants to select those located within10 kb of these genes (GENCODE v36 annotations^97^; 2,954 variants) or within known COVID-19-associated loci (17 variants), defined as variants in LD (r^2^>0.8 in 1KGP Europeans) with lead variants from the *B2_ALL_eur_leave_23andme* GWAS (release 4, COVID-19 Host Genetics Initiative^98^).

To capture distant regulatory effects, we extended this set to include variants overlapping putative distal enhancers that physically interact with immune-relevant genes in immune cells or lung tissue^99^ (1,190 variants). Using promoter capture-HiC data^52^, we identified putative distal enhancers active in immune cells (CHICAGO score >5), retrieving 983 variants overlapping a candidate regulatory element associated with at least one immune-relevant gene in myeloid (Mon, Mac0, Mac1, Mac2) or T cell (nCD4, tCD4, aCD4, naCD4, nCD8, tCD8) contexts. To account for regulatory activity in the lung − a primary-interface with environmental pathogens − we incorporated promoter capture-HiC data from lung tissue^53^, identifying 207 additional variants within putative distal enhancers interacting with immune-relevant genes. This yielded a total of 4,161 archaic variants to be included in the MPRA library.

#### Positive and scrambled controls

As positive controls for regulatory activity, we selected 113 fragments from putative promoter sequences, defined from CAGE-seq data^100^.

Specifically, we downloaded CAGE-seq data from fantom5 phase 2 (https://fantom.gsc.riken.jp/5/datafiles/phase2.0/extra/CAGE_peaks/hg19.cage_peak_phase1and2combined_tpm_ann.osc.txt.gz) and selected the top 40 most active TSS from three distinct tissues: resting CD14+ monocytes (n=9 samples), CD4 and CD8 T cells (n=38 samples) and airway epithelial cells (alveolar, bronchial, small airways, and tracheal; n=20 samples), corresponding to a list of 93 unique promoters, to which we added 20 additional promoters selected as the monocyte TSS showing the strongest fold changes in response to stimulation with either LPS or IFN (n=6 samples). We then extracted the 270 bp sequence around the center of each promoter as positive control fragments. Positive control sequences were then shuffled at random and included in the library to serve as “scrambled” controls for cCRE activity.

### Lentivirus-based MPRA experiments

#### Oligonucleotide pool design

For each of the 4,274 cCREs in the final dataset, two 270-bp DNA fragments were designed, carrying either the introgressed or the non-introgressed allele (or promoter/scrambled version for the control cCREs). All fragments were based on the GRCh37 human reference genome, with the focal allele positioned at the center of the sequence. For each putative regulatory sequence, the strand used for oligonucleotide design was selected according to the orientation of the nearest gene (when located within 10kb) or the predicted target gene (for promoter-interacting regions, PIRs). When a PIR was associated with genes located on both strands, the strand with the largest number of genes was prioritized. Fragments containing homopolymer tracts longer than 10 bp were excluded to minimize errors during oligonucleotide synthesis and PCR. For 107 variants in which such homopolymers were located within 50 bp of either fragment end, alternative homopolymer-free sequences were designed by shifting the position of the focal allele by 50 bp upstream (n=55) or downstream (n=52) from the center. Finally, 15-bp adapter sequences were added to both ends of each fragment (5’-AGGACCGGATCAACT-3’ and 5’-CATTGCGTGAACCGA-3’) to enable downstream amplification and cloning, resulting in a total fragment length of 300 bp. All sequences were synthesized as single-stranded oligonucleotides by TWIST Bioscience.

#### Generation of lentiMPRA libraries and sequencing

The oligonucleotide pool was first amplified by 14 cycles of PCR using Kapa PCR mix (Kapa Biosystems) with primers complementary to the adaptor sequences flanking each oligonucleotide (forward: 5’-AGGACCGGATCAACT-3’; reverse: 5’-TCGGTTCACGCAATG-3’). The lentiMPRA library was generated as previously described^40^. Briefly, a minimal promoter (mP) was added downstream of each oligonucleotide by a five-cycle PCR (primers 5BC-AG-f01 and 5BC-AG-r01), with NEBNext Ultra II Q5 Master Mix, followed by an additional eight-cycle PCR to introduce a 15-bp random barcode downstream of the mP (primers 5BC-AG-f02 and 5BC-AG-r02) (Supplementary Table 7). The resulting oligo-mP-barcode pool was cloned into the pLS-SceI vector (Addgene, 137725) by Gibson assembly recombination using NEBuilder HiFi DNA assembly Master Mix (NEB). The purified recombinant product was electroporated into 10-beta competent cells (NEB) using a GenePulser Xcell electroporation system (Bio-Rad; 2 kV, 25 uF, 200 ohms). Colonies grown overnight on 10cm LB agar plates with 10 μL of 100 mg/mL carbenicillin were pooled, and plasmid DNA was extracted using the EndoFree Plasmid Maxi kit (Qiagen). Two independent plasmid libraries were generated. For library 1, colonies were collected to achieve an average of ~200 barcodes per oligonucleotide, whereas library 2 targeted ~100 barcodes per oligonucleotide. For the second library, PCR conditions were optimized to improve the representation of low-GC oligonucleotides (Supplementary Fig. 1).

To map barcodes to their corresponding cCRS, each library was sequenced as previously described^40^. Briefly, fragments containing the cCRS, mP, and barcode were amplified and sequenced: library 1 with four MiSeq runs (2×250bp) using custom primers (R1, pLSmP-ass-seq-R1; R2[index read], pLSmP-ass-seq-ind1; R3, pLSmP-ass-seq-R2; Supplementary Table 7), yielding ~56 million reads, while library 2 with a single Nextseq 2000 run (2×-150 bp) to a depth of ~58 million reads.

#### Cell culture, lentivirus packaging and titration

Human lung epithelial A549-derived ACE2plusC3 cells (ATCC, CRL-3560) and hepatocyte HepG2 cells were cultured in DMEM supplemented with 10% heat-inactivated fetal bovine serum (FBS). Hematopoietic progenitor K562 cells were cultured in RPMI 1640 supplemented with 10% heat-inactivated FBS. All cell lines used were tested negative for mycoplasma. Lentivirus carrying the lentiMPRA library (hereafter referred to as lentivirus) was generated by co-transfecting the lentiMPRA plasmid library with psPAX2 and pMD2.G in nine T150 flasks of HEK293T cells using Fugene HD (Roche) or EndoFectin (Genecopoeia). After 8-9 h, cell culture media was refreshed and supplemented with Viralboost (Alstem). The produced lentivirus in the conditional medium was collected 2 days after transfection, filtered through a 0.45 mm PES filter system (Thermo Fisher Scientific) and concentrated using Lenti-X concentrator (Takara Bio). Lentiviral titration was performed as previously described^40^. Briefly, lentivirus was titrated on each cell type using a serial of volumes (0, 0.5, 1, 2, 4, 8, 16, 32μl) along with 8μg/ml of Polybrene (Sigma-Aldrich, TR-1003). Viral genome copies were quantified by qPCR (Supplementary Table 7) using Power SYBR™ Green PCR Master Mix (Applied Biosystems™, Cat No. 4367659), following manufacturer’s instructions, to estimate the Multiplicity of infection (MOI). Based on virus titer and infection efficiency, the optimal cell number and lentivirus input volume were determined for each cell type.

#### Lentiviral infections, stimulations, and barcode sequencing

Approximately 6 million A549plusC3 and HepG2 cells, and 10 million K562 cells, were infected per replicate and condition. A549plus-C3 and HepG2 cells were seeded into 2-3 Petri dishes-100, and K562 cells were seeded in one T150 flask per replicate and condition. Cells were infected with lentivirus in the presence of 8 μg/mL polybrene to increase infection efficiency. Polybrene was removed 24 h post-infection by refreshing culture media. Cells were infected at an estimated MOI of 60, 90, and 45 for A549plus-C3, HepG2, and K562, respectively, and cultured for 48 h. After the 48h-lentivirus infection, cells were treated for 6 h with various stimuli: ACE2plusC3 cells with IAV (H1N1 influenza virus strain A/PR8/1934, MOI=1), SARS-Cov-2 (GE1973/, MOI=1), produced as previously described^29^, or TNF-α (10 ng/ml); and HepG2 and K562 cells with IFN-α2b (500pM, 10 ng/ml), dexamethasone (1 μM, 400ng/ml), or TNF-α (10 ng/ml). Recombinant IFN-α2b was provided by D. Gewert (Wellcome Foundation, Beckenham, Kent, UK; now at BioLauncher Ltd, Cambridge, UK). Dexamethasone was obtained from Sigma (catalog n° D4902) and TNF-α was purchased from Proteintech Group, Inc (catalog n° HZ-1014). Three to five replicates were performed per cell type for each stimulation condition, ensuring that every stimulated replicate had at least one unstimulated replicate from the corresponding cell line, performed from the same lentivirus preparation.

Following stimulation, cells were washed and lysed in RTL buffer and stored at −80°C until RNA and DNA extraction using the AllPrep mini kit (Qiagen). RNA samples were treated with DNase using both on-column digestion (RNase-Free DNase Set, Qiagen) and in-solution treatment (TURBO DNA-free Kit, Qiagen). cDNA was synthesized from total RNA with SuperScript II RT (Invitrogen) with a primer containing a unique molecular identifier (UMI) (P7-pLSmp-ass16UMI-gfp, Supplementary Table 7). DNA fragments were amplified from both genomic DNA and cDNA, keeping replicates and nucleic acid types separate, using a three-cycle PCR with primers containing sequencing adapters (P7-pLSmp-ass16UMI-gfp and P5-pLSmP-5bc-i#, Supplementary Table 7). These primers included sample-specific indexes for demultiplexing and UMIs for deduplication. A second PCR round was used to amplify the library for sequencing with primers targeting the adapters (P5 and P7). Fragments were pooled and purified as previously described^40^. DNA and RNA fragments were combined across conditions and sequenced on HiSeq X or NextSeq2000-C50 platforms, achieving an average depth of >10 reads per DNA barcode and >30 reads per RNA barcode, using custom sequencing primers (Supplementary Table 7).

### MPRA processing pipeline

#### Barcode-cCRS association dictionary

Barcode-cCRS association was performed using a modified version of the *mpraflow pipeline*^40^. Specifically, paired-end 270-bp reads from the lentiMPRA plasmid library were merged using *fastq-join* version 1.3.1 (ref.^101^) and aligned to the reference sequences with *bwa* version 0.7.17 (ref.^102^). To accurately quantify the activity of SNVs, we excluded any read that did not match the 270-bp reference sequence of its associated cCRS, minimizing artifacts in barcode-cCRS associations that persisted even when using the *--cigar 270M* option. Filtered reads were then used to assign barcodes to cCRS, requiring a minimum of three supporting reads per barcode-cCRS association and excluding barcodes mapping to more than one cCRS. To further reduce misassignment, barcodes with sequence complexity <11 were removed, where sequence complexity was defined as the Shannon entropy of nucleotide frequencies within the barcode:

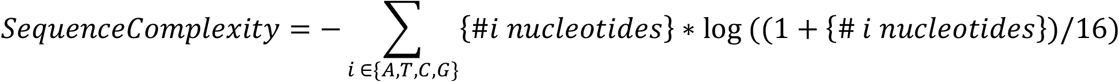

This threshold excludes fewer than 0.1% of truly random 15-bp sequences while removing sequences with highly skewed nucleotide composition. Using this procedure, we recovered 95.2% and 96.6% of tested cCRS per library, of which 90.5% and 97.1% had >10 barcodes, yielding a median of 108 and 80 barcodes per oligo for libraries 1 and 2, respectively.

#### Transcription rate quantification

Fastq files for each DNA and RNA samples were imported using the *readDNAStringSet* function from the *BioStrings* R package^103^. We excluded mismatched read pairs where barcode sequences did not perfectly match between the two insert reads (R1 and R2), unassigned barcodes absent from the corresponding plasmid library dictionary, and duplicated UMIs associated with more than one barcode from the same library. For each replicate sample, we counted the number of RNA and DNA UMIs per barcode and estimated size factors to reflect the complexity of each cDNA library, separately for RNA and DNA UMIs and for each plasmid pool. Size factor estimation was performed as in *DEseq*^104^. Specifically, for each cDNA library *j* corresponding to nucleic acid type *n* (RNA or DNA) of cells transfected with a lentivirus library derived from plasmid pool *p*, the size factor *s*_*j*,_^(*n*,*p*)^ was computed as:

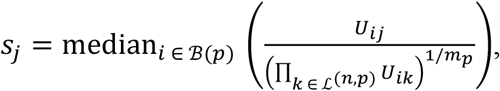

where *U*_*ij*,_ is the number of UMIs per million associated with barcode *i* and cDNA library *j*. ℬ (*p*), the set of barcodes associated with a known cCRS in plasmid pool *p*, and ℒ^(*n*,*p*)^ the set of all cDNA libraries of nucleic acid type *n* derived from plasmid pool *p*, of size *m*_*p*_. The size-adjusted depth of each cDNA library *j* was computed as the product of the total number of UMIs and its estimated size factor.

For each barcode *i* associated to cCRS *j*, an outlier score was calculated

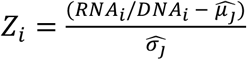

where 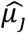 and 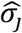 are the winsorized mean and standard deviation of RNA/DNA ratios across all barcodes for cCRS *j*. Barcodes with |*Z*_*i*_| >2 were excluded prior to transcriptional activity quantification.

To aggregate the barcodes and quantify the transcription rate induced by each cCRS, we first applied *analyzeQuantification* from the *MPRAnalyze* framework^57^ separately to each cCRS, condition and replicate, estimating cCRS activity with *getAlpha*(). In addition, for each cell type and stimulation condition, DNA and RNA counts were jointly modeled across replicates to obtain average oligo activity per cell type and condition. Because cCRS activity correlated with GC content, even among scrambled sequenced (Spearman’s ρ>0.22, *P*<0.03; Supplementary Note 3), activities were adjusted for GC content within each cell type, by fitting a linear model on scrambled sequences and subtracting the estimated effect of GC 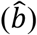 across all cCRS. Specifically, GC effect was estimated on the log scale:

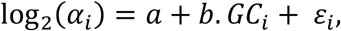

where *GC*_*i*_ is the GC content of *i*^*th*^ *cCRS, α*_*i*_ its activity in the current cell type and condition, *b* the estimated GC effect, *a* the intercept, and ε_*i*_ the normally distributed residuals. Corrected activities were computed as:

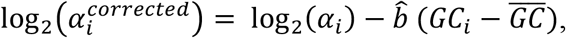

where 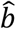 is the GC effect from the scrambled sequences, and 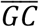 is the mean GC content of tested cCRS. Within each cell type and condition, GC-corrected activities were centered on the log_2_ scale so that the median RNA/DNA ratio of all tested sequences equaled 1. Log-activities and their standard errors were further scaled by the mean absolute deviation to ensure consistent variance across cell types and conditions.

#### Variance decomposition analysis and UMAP

To estimate the proportion of variance explained by cell type and stimulation condition, we used GC-corrected, normalized activity estimates per replicate and fitted, for each cCRS, a linear mixed model (LMM) of the form:

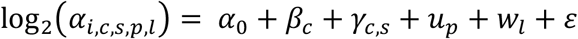

where *α*_*i*,*c*,*s*,*p*,*l*_ denotes the cCRS activity of replicate *i* from cell type *c* under stimulation condition *s*, derived from plasmid library *p* and lentivirus preparation *l*. The term 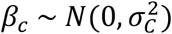 captures the effect cell type on activity, 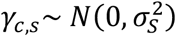 the cell type-specific effect of stimulation, 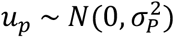 the effect of plasmid library, 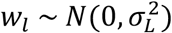 the effect of lentivirus preparation, and 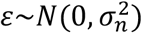 residual noise between replicates. We then computed,for each cCRS, the proportion of variance explained by cell type g _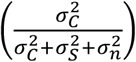_ and by stimulation condition 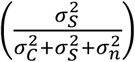. Batch-corrected activities were defined as 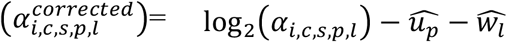, where 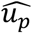 and 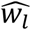 are derived from the LMM using the *ranef ()* function. Finally, UMAP was performed on batch-corrected activities using the *umap()* function of the *umap* R package^105^ with option min.dist=0.5.

#### Detection of active cCREs

To evaluate whether tested cCRS significant modulated expression of a downstream minimal promoter, we used the *analyzeComparative* function from the *MPRAnalyze* package with option *fit*.*se=TRUE* to estimate standard errors associated with activity estimates. Analyses were performed separately for each cell type and condition, modeling DNA and RNA counts jointly across replicates, with identical full model and reduced model. The standard error associated with the intercept was extracted as a measure of uncertainty in log-transformed activity estimates. Activity Z-values were computed for each oligo by dividing GC-corrected log-transformed activity by its corresponding standard error. To validate this procedure, we implemented a permutation strategy to generate mock oligos mimicking the behavior of inactive sequences (Supplementary Fig. 3a). Specifically, barcodes were randomly reassigned across cCRS to (i) preserve the distribution of the number of barcodes per cCRS as in the true data set and (ii) set the expected activity of each cCRS to the mean activity across all tested sequences (null activity). Transcription rate quantification was repeated on this permuted dataset, confirming that, in the absence of regulatory activity, Z-scores follow a standard gaussian distribution (Supplementary Fig. 3b).

For each cell type and condition, we compared normalized activity values between observed and permuted datasets to compute empirical *p*-values and estimate FDR. For any cCRS with absolute *Z*-score |Z_i_| in the observed data, the empirical *p*-value was defined as the proportion of permuted sequences with an absolute Z-score > |Z_i_|. At a given *Z*-score threshold 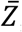, the FDR was estimated as the ratio of the number of cCRS with an absolute Z-value 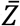, between the permuted and the observed dataset.

cCRS with *FDR<0*.*05, and* 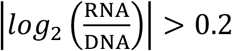 were considered as active. Active cCRS were further categorized as having upregulating 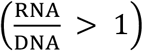 or downregulating 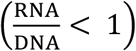 activity. In addition, active cCRS with 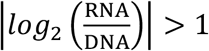 were classified as strong, whereas those with 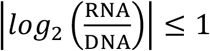 were classified as weak. For cCREs where transcriptional activity differed between the two cCRS, the cCRE classification was assigned based on the cCRS with the strongest activity, following the hierarchy: strong upregulating cCRE> weak upregulating cCRE > strong downregulating cCRE> weak downregulating cCRE > inactive.

#### Enrichment of active cCREs in chromatin states and TFBM

For each cell type, cCREs were assigned to one of 15 chromHMM states using ENCODE annotations^54^ and classified as strong/weak upregulating cCRE or downregulating cCRE based on their transcriptional activity in unstimulated cells. Odds ratios were computed to quantify the overlap between cCRE activity classes and chromHMM across the three cell types. For TFBM enrichment, we retrieved position frequency matrices for all human TFs from JASPAR Core 2020 (ref.^56^).

Binding affinities were computed using the *searchSeq* function from *TFBStools*^106^ (*min*.*score*= “0%”, *strand* = “*”), yielding log-transformed binding affinity scores at each position along the sequence. After reverting to the natural scale, we summed, for each TF and sequence, the binding affinities across all positions and both strands to obtain a single activity measure, which was then log-transformed again, as described previously^107^. These values were then adjusted for total sequence GC content. For each TF, the top 10% of sequences with the highest GC-adjusted binding affinity were defined as putative targets. Enrichment of active cCREs (split by transcriptional activity in unstimulated cells) within these was assessed using Fisher’s exact test. A cCRE was considered bound by a TF if at least one of its two alleles was classified as a putative target.

#### Detection of cell type-specific and stimulation-responsive cCREs

To identify sequences with differential transcriptional activity across cell types and stimulation conditions, we modeled RNA and DNA counts using the *analyzeComparative* function from the *MPRAnalyze* package with the option *fit*.*se=TRUE*. Library depth for each sample was defined as the total number of UMIs multiplied by the corresponding size factor.

To assess cell type-specific effects, analyses were restricted to the unstimulated condition and used the following design for each pair of cell types:

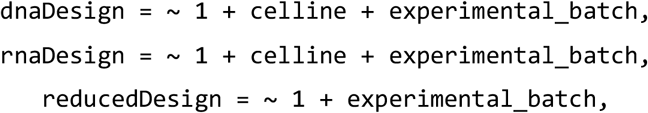

where *experimental_batch* corresponds to the combination of plasmid library and lentivirus preparation batch. For responsive cCREs, we compared stimulated and unstimulated replicates within each cell type using:

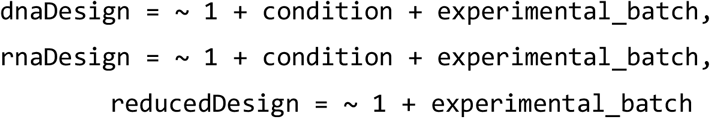

In both cases, *p*-values for differential activity were computed using *testLrt* function. Standard errors of the estimated logFC were extracted from the *r*.*se* element of the *modelFits* object. LogFC returned by testLrt and their standard errors were divided by log(2) to obtain log_2_ fold changes in activity. To validate the calibration of *p*-values and control the FDRs, we repeated all analyses on permuted data in which sample labels were randomly assigned across barcodes (Supplementary Fig. 5), ensuring that each cCRE had identical activity across cell types or conditions. For each comparison, empirical *p*-values and FDR were estimated as described for activity. Differences in TF binding affinity between responsive and non-responsive cCREs were assessed using a Wilcoxon rank-sum test.

#### Detection of emVars and context dependency

To identify expression-modulating variants (emVars), we jointly modeled RNA and DNA counts from both alleles using the *analyzeComparative* function of the *MPRAnalyze* package (*option fit*.*se=TRUE*), setting size factor-normalized total UMI counts as library depth. EmVars were assessed separately for each cell type and condition using the following design:

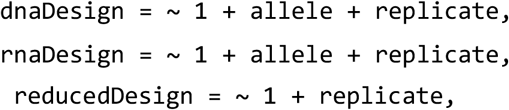

*P*-values for differential activity between alleles were computed using the *testLrt* function. Standard errors of the estimated logFC were extracted from the *r*.*se* element of the *modelFits* object. LogFC returned by testLrt and their standard errors were divided by log(2) to obtain log_2_ fold change in activity. To validate *p-*value calibration and control FDRs, we repeated the analyses on permuted data in which labels were randomly reassigned across barcodes, ensuring identical activity between alleles for each cCRE (Supplementary Fig. 6). Empirical *p*-values and FDR were computed as described above. To limit multiple testing and increase power, emVar discovery was restricted to variants and conditions where at least one allele had been classified as active. No activity filter was used when replicating emVars across conditions.

To identify context-dependent emVars, we extended this framework by jointly modelling activity across conditions and testing for allele-by-condition interactions. For cell type-specific emVars, analyses were restricted to the resting condition and used the following design for each pair of cell types:

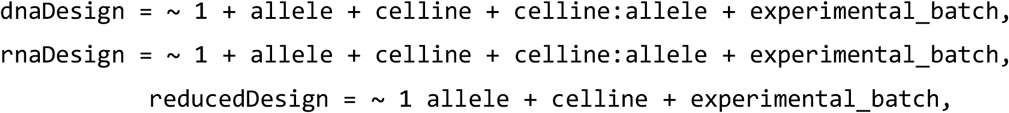

where *experimental_batch* corresponds to the combination of plasmid library and lentivirus preparation batch. For stimulation-responsive emVars, we compared allelic effects between unstimulated and stimulated conditions within each cell type using:

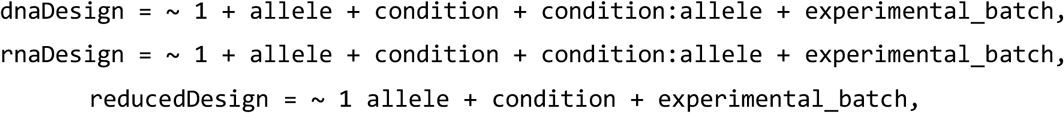

In both cases, interaction *p*-values were computed using *testLrt* function. Standard errors of the estimated logFC were extracted from the *r*.*se* element of the *modelFits* object. LogFC returned by testLrt and their standard errors were divided by log(2) to obtain log_2_ fold change in allelic effects. For context-dependent emVars, permutations were performed as in the differential activity analyses by shuffling condition labels across barcodes, such that allelic effects remained constant across cell types or conditions. Empirical *p*-values were computed as described above. For FDR estimation, only interactions involving variants previously identified as significant emVars in at least one of the two contexts compared (for example, resting or stimulated) were considered.

To assess impact of responsive emVars on TFBMs, sequences overlapping each variant were scanned using FIMO^108^ and the JASPAR Core 2020 database^56^. For each locus, TFs with significant motifs matches and the largest predicted differences in binding affinity between the two alleles were reported.

#### Comparisons with external datasets

To assess the reproducibility of emVars, we retrieved detected emVars from a previous plasmid-based MPRA study^49^ in which 22,412 variants were tested across five different cell types, including HepG2 and K562. We identified 160 variants shared between the two studies and compared effect sizes for emVars detected in at least one dataset. To assess the overlap between emVars and molecular QTLs and GWAS traits, we downloaded 95% credible sets for these associations from the Open Targets platform^109^ (http://ftp.ebi.ac.uk/pub/databases/opentargets/platform/25.03/output/credible_set), and extracted all traits for which the 95% credible set contained at least one of the 3,838 archaic variants tested in our MPRA. This yielded 1,810 eQTL-variants (eQTLtype = ‘ge’) associated with 4,594 expression traits; 1,668 splice or transcript usage QTL-variants (studyType==‘sQTL’ or ‘tuQTL’) associated with 5,515 splicing traits; and 1,745 GWAS-variants (studyType==‘gwas’) associated with 2,045 phenotypic traits. Concordance of effect direction between emVars and eQTLs was assessed by comparing the unstimulated emVar effect sizes with the effect sizes associated with the introgressed allele in each QTL study. For each emVar, effects were classified as fully concordant if the direction was concordant across all tissues or studies in which the emVar overlapped a significant QTL, partially concordant if concordant in at least one tissue or study, and fully discordant if the direction was opposite across all tissues or studies.

#### Local emVar enrichments

To identify specific expression phenotypes underlying adaptive introgression in humans, we tested each introgressed region for local enrichment of regulatory alleles with consistent directional effects under specific conditions. For each introgressed locus containing more than five tested alleles, we counted the number of alleles that significantly increased (P_emp_<0.05, log_2_FC>0), or decreased (P_emp_<0.05, log_2_FC<0) reporter gene expression. Enrichment was assessed used Fisher’s exact test, comparing observed counts to genome-wide expectation, with separate tests for expression-increasing and expression-decreasing alleles. Multiple testing correction was performed using the Benjamini-Hochberg procedure across all loci and conditions, separately for expression-increasing and expression-decreasing alleles. This enrichment analysis was also performed at the gene level, considering 10-kb windows centered on each gene.

#### Comparison of emVar occurrence across populations, and archaic sources

To assess the occurrence of emVars across conditions and among specific subsets of introgressed alleles, we partitioned introgressed alleles by inferred archaic source and recipient population. For each group and condition, the proportion of non-null hypothesis *π*_1_ among differential activity tests was estimated. Specifically, given that empirical *p*-values are uniformly distributed under the null, *π*_1_ can be estimated for a given significance threshold *α* (with sufficient power) as:

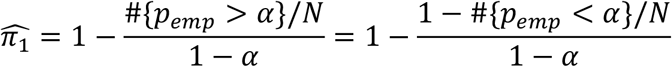

where *N* is the total number of tests performed for the group and condition considered. In practice, *α* = 0.05 was used.

To estimate the proportion of alleles associated with increased or decreased expression, this framework was extended as follows:

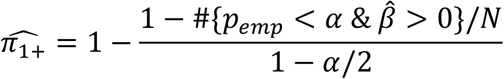

and

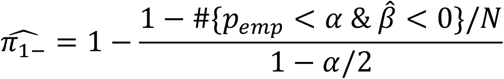

For each group and condition, 95^%^ confidence intervals were estimated using 1,000 bootstrap replicates (resampling variants at random within each group, with replacement). Differences in proportion of regulatory alleles between archaic sources were assessed using Fisher’s exact test, comparing the proportion of variants with *p*_*emp*_ < 0.05, between Neanderthal- and Denisova-inherited variants.

### CRISPR activation

To identify the target gene regulated by the rs17713054 enhancer, we generated A549 cells stably expressing the nuclease-deficient Cas9 fused to transcriptional activation domains (dCas9-VPR) by lentiviral transduction. Lentiviral particles were produced using the Edit-R CRISPRa Lentiviral hCMV-Blast-dCas9-VPR Plasmid (Horizon Discovery, CAS11914). A total of 1 x 10^5^ cells from two independent Blasticidin-resistant A549 pools were seeded in 24-well plates and transfected with either 7.5 pmol of a pool of three single guide RNAs (sgRNAs), a single sgRNA targeting the rs17713054 enhancer, or a negative control sgRNA (Gal4-4, GAACGACTAGTTAGGCGTGTA)^110^. Transfections were performed using Lipofectamine RNAiMAX (Life Technologies, catalogue n° 13778). Total RNA was extracted 6 or 24 h after transfection using the PureLink® RNA Mini Kit (Life Technologies, catalogue n° 12183020) with on-column DNase digestion (Rnase-free Dnase Set, Qiagen, catalogue n° 79256). cDNA was synthesized using SuperScript II reverse transcriptase (Life Technologies, catalogue n° 18064-071). Gene expression levels of *TBP* (used as housekeeping gene), *SLC6A20, LZTFL1*, and *CXCR6* were measured by qPCR using Power SYBR™ Green PCR Master Mix (Applied Biosystems™, catalogue n° 4367659), with gene-specific primers (Supplementary Table 7).

To investigate the genome-wide effect of CRISPRa on gene expression profiles, poly(A)-enriched mRNA libraries were prepared from 1,000 ng of total RNA using the Illumina® Stranded mRNA Prep kit (catalogue n° 20040534) from cells transfected with the rs17713054-targeting sgRNA or negative control sgRNA (n=6 biological replicates).Libraries were sequenced on a NextSeq 2000-100C platform (60-bp paired-end reads). Reads were aligned to the GRCh38 reference genome and quantified using the Sequana RNA-seq pipeline^111^, with the *sequana_rnaseq* command with option *--aligner-choice star*, and gene annotations from *Ensembl GRCH38 v113*. Differential expression analysis was performed using the *rnadiff* command^111^, setting *--minimum-mean-reads-per-condition-per-gene 5* and adjusting for replicates effects.

## Data availability

All sequences data generated in this study, including barcode-cCRS association libraries and UMI count data, have been deposited in the European Nucleotide Archive (accession n° XXX). Genome data from Choin *et al*. were retrieved from the European Genome-Phenome Archive (EGA, accession code EGAS00001004540). SGDP genome data were retrieved from the European Nucleotide Archive (ENA, accession codes PRJEB9586 and ERP010710). Genome data from Malaspinas *et al*. were retrieved from the EGA (accession code EGAS00001001247). Genome data from Vernot *et al*. were retrieved from dbGAP (accession code phs001085.v1.p1).

## Code availability

All code used for MPRA processing and data analyses presented in this study can be accessed at https://github.com/h-e-g/MPRA_archaic_immunity.git

## Acknowledgments

The authors thank Aleksi Isomursu (Institut Pasteur) for helpful discussions on CRISPR activation, and Marc Monot, Laurence Ma, and Etienne Kornobis (Biomics Platform, Institut Pasteur) for their contribution to NGS sequencing and RNA-seq analyses. The Human Evolutionary Genetics Unit is supported by the Institut Pasteur; the Collège de France; the Centre Nationale de la Recherche Scientifique (CNRS); the Agence Nationale de la Recherche (ANR) grants POPCELL-REG (ANR-22-CE12-0030-01), COVIFERON (ANR-21-RHUS-08), and France 2030-EPIGEMI (ANR-23-CHBS-0007); the European Union’s HORIZON-HLTH-2021-DISEASE-04-07 grant UNDINE (101057100); the French government’s Investissement d’Avenir program ‘Laboratoires d’Excellence’ Integrative Biology of Emerging Infectious Diseases (ANR-10-LABX-62-IBEID) and Milieu Intérieur (ANR-10-LABX-69-01) projects. The Biomics Platform (Institut Pasteur) is supported by France Génomique (ANR-10-INBS-09) and IBISA.

## Author contributions

Z.L, L.Q.-M. and M.R. planned and designed the study. Z.L. performed all MPRA and CRISPRa experiments. M.R. performed all bioinformatic analyses, with contributions from G.K and J.M.-R. G.K., J.M.-R. and E.P. provided expertise in archaic introgression analyses, and F.I., S.F. and D.G. provided specialist expertise in MPRA design. Z.L, L.Q.-M. and M.R. wrote the manuscript, with input from G.K., J.M.-R., F.I., S.F., E.P. and D.G.

---

**Extended Data Fig. 1.**
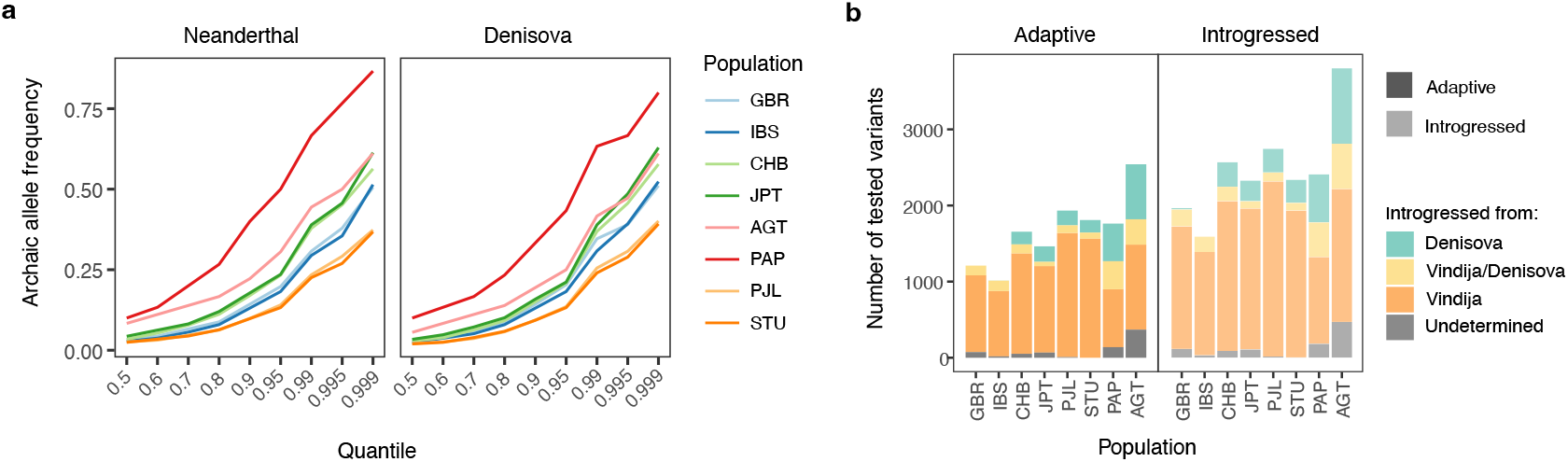
Frequency of introgressed variants across populations and geographic distribution of tested introgressed alleles. **a**, Quantiles of the frequency of archaic-derived alleles across the eight studied populations, shown separately for alleles matching either the Vindija Neanderthal or the Denisovan genomes. **b**, Number of archaic variants tested per population, colored according to the inferred source of introgression. Left: considering only putatively adaptive alleles in each population (frequency above the 95^th^ percentile). Right: considering all common variants. Note that the same variant may be counted multiple times across different populations.

**Extended Data Fig. 2.**
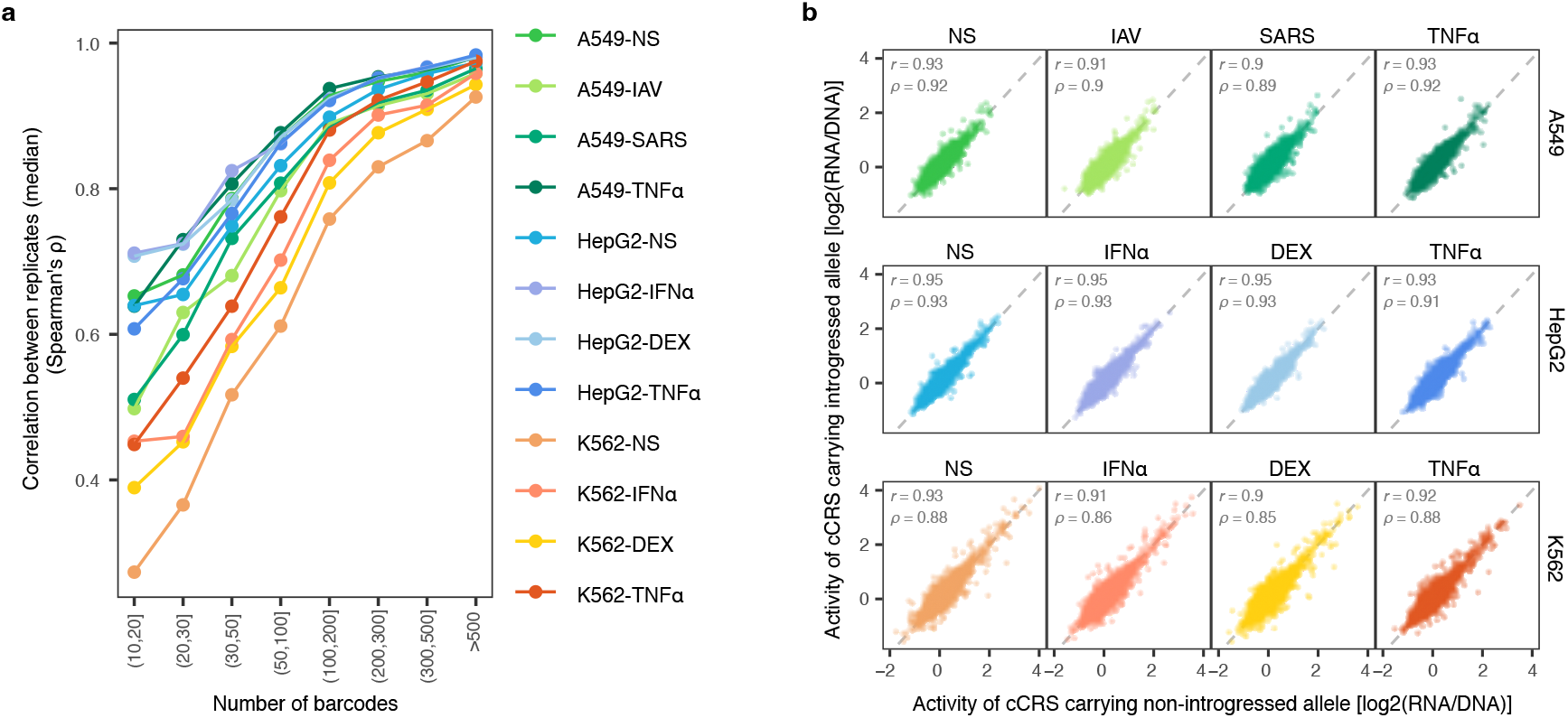
cCRE activity shows strong correlation between alleles and replicates. **a**, Median correlation of cCRS activity between replicates, shown as a function of the average number of associated barcodes across libraries 1 and 2. Each line represents a specific cell type and stimulation condition. **b**, Correlation of cCRS activity between introgressed and non-introgressed alleles across all tested conditions. Pearson’s *r* and Spearman’s ρ are reported for each condition.

**Extended Data Fig. 3.**
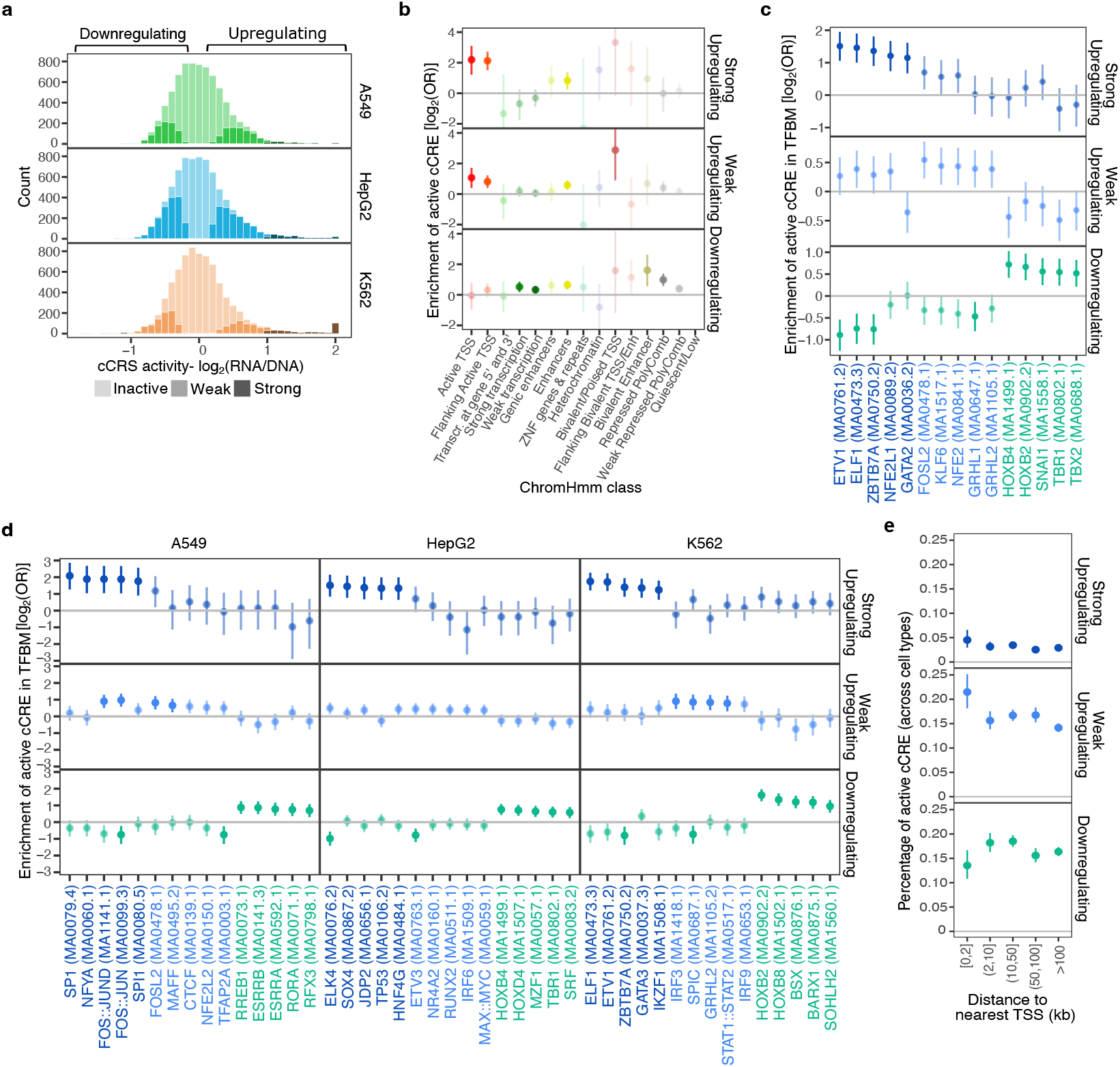
Upregulating and downregulating cCREs display specific epigenetic marks and transcription factor binding motifs. **a**, Distribution of log_2_-transformed cCRS activity across three cell types (unstimulated condition only). Bright colors denote active cCRSs (FDR<5%), with darker colors indicating strongly upregulating cCREs (log_2_(DNA/RNA) >1). **b**, Enrichment of active cCREs across chromatin marks defined by ChromHMM. **c**,**d**, Enrichment of active cCREs in transcription factor binding motifs (TFBMs), analyzed either jointly across all three cell types (**c**) or separately in each cell type (**d**). **e**, Proportion of cCREs assigned to each transcriptional activity class as a function of distance to the nearest active transcription start site (TSS), defined using ENCODE CAGE-seq data. **b-e**, Vertical bars indicate 95% confidence intervals. In **b-d**, enrichments not reaching the 5% FDR threshold are shown as transparent. In **c**,**d**, for each category of active cCREs, only 5 TFs with the highest lower bound of the 95% confidence interval are shown, with a maximum of two TFs per family.

**Extended Data Fig. 4.**
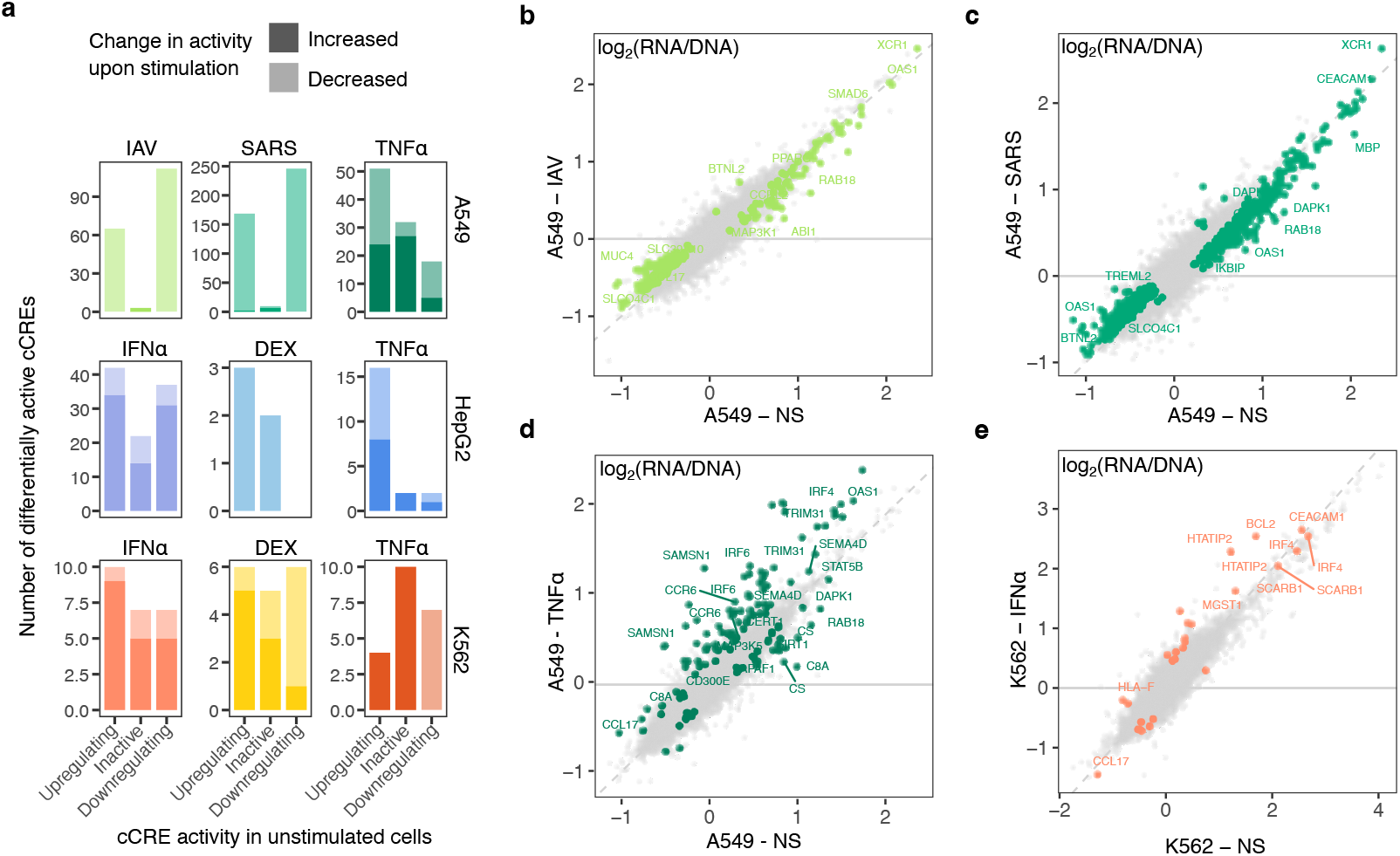
Impact of stimulation on cCRE activity. **a**, Total number of cCREs showing activity changes upon stimulation, stratified by cCRE class in the unstimulated condition. Color indicates the direction of change, darker shades denoting increased activity and lighter shades decreased activity. For downregulating cCREs, increased activity corresponds to reduced transcription. **b-e**, Comparison of transcriptional effects of tested sequences (one dot=1 cCRS) between unstimulated and stimulated conditions. Only the four conditions with the highest number of responsive cCREs are shown.

**Extended Data Fig. 5.**
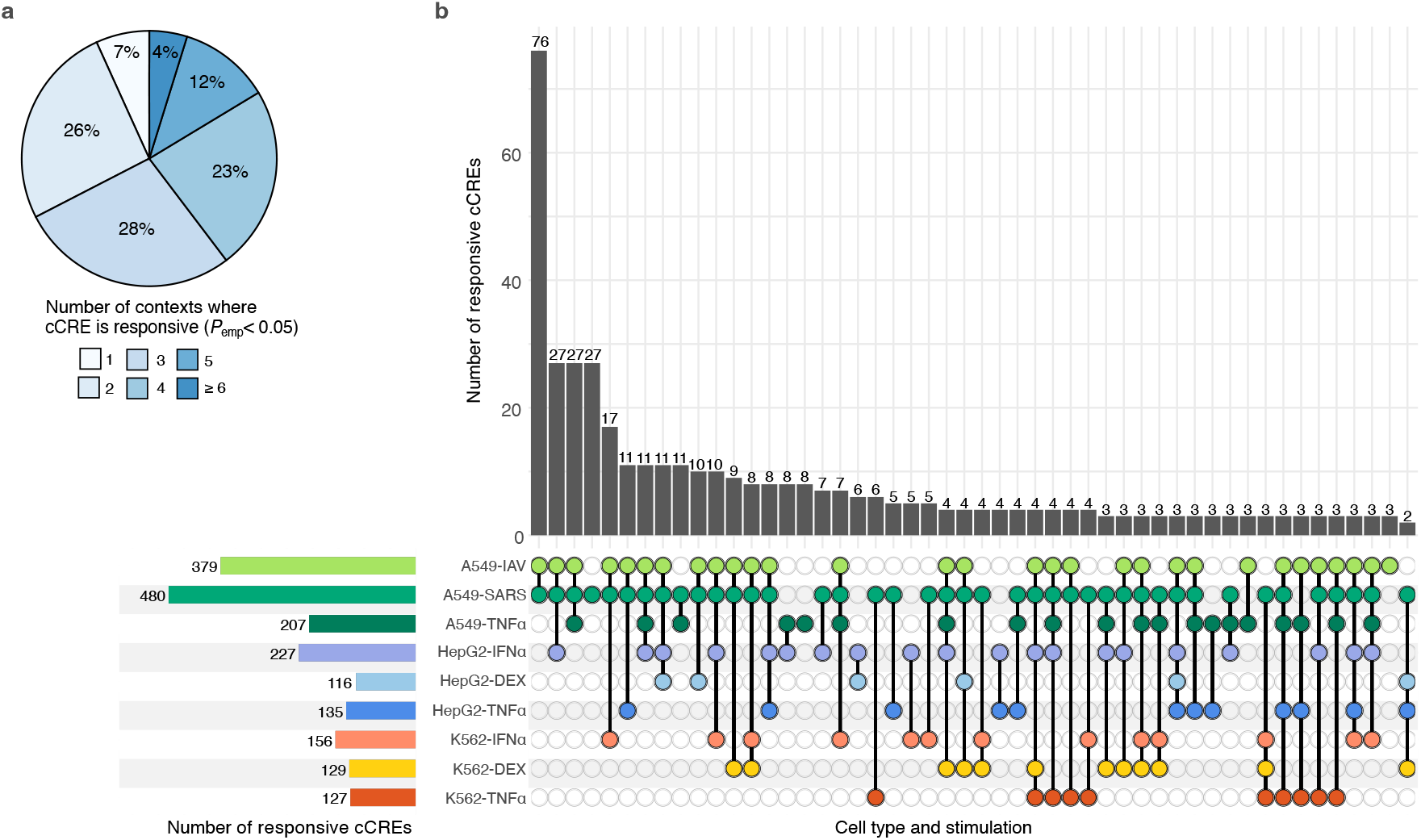
Sharing of responsive cCREs (relaxed threshold). **a**, Proportion of responsive cCREs (FDR<5% for discovery) that are context-specific or shared across multiple contexts when using a relaxed *p*-value threshold for replication (*P*_*emp*_<0.05). **b**, UpSet plots showing the frequency of responsive cCREs across contexts (cell types × stimuli, bottom left) and across specific combinations of contexts (right), using FDR<5% for discovery and relaxed *p*-value threshold (*P*_*emp*_<0.05) for replication. Only combinations with two or more responsive cCREs are shown.

**Extended Data Fig. 6.**
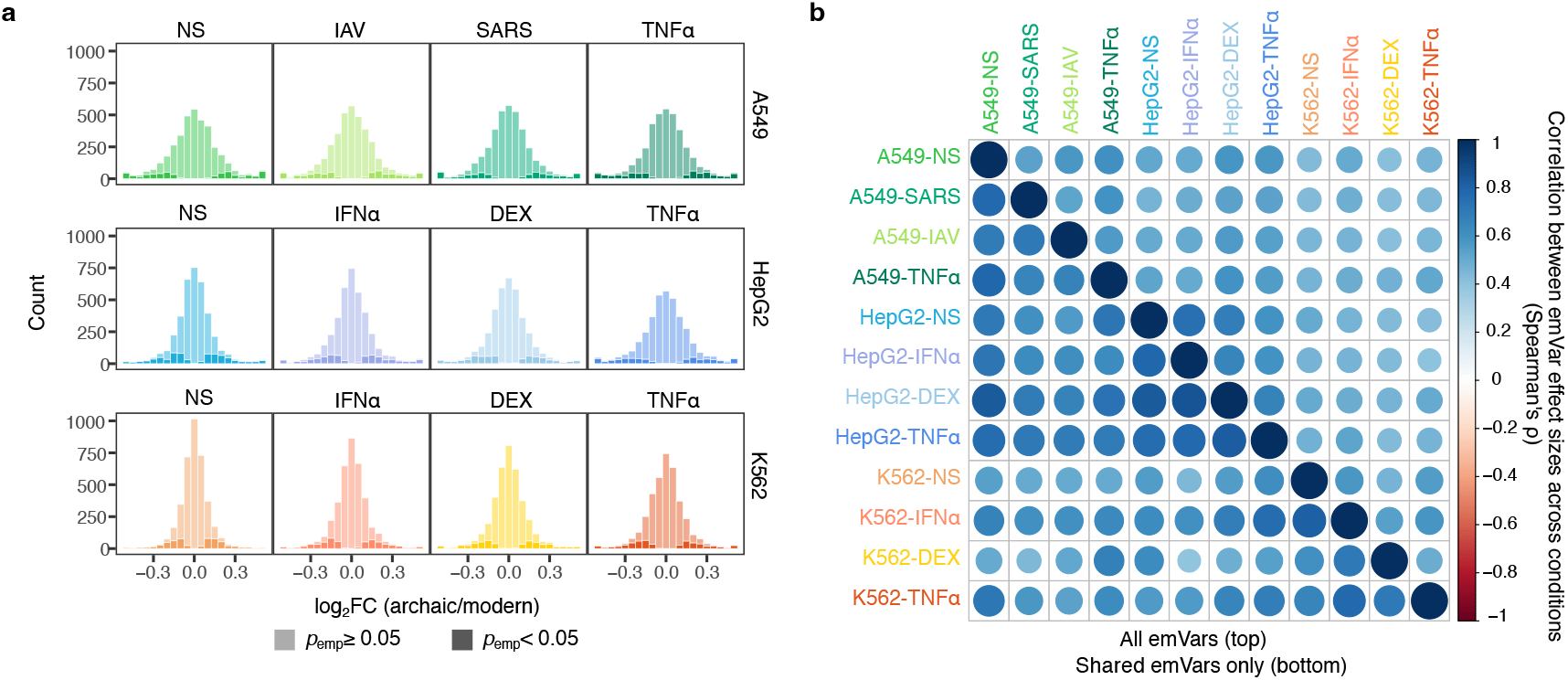
Distribution of emVar effect sizes and sharing across conditions. **a**, Distribution of estimated effect sizes for all tested emVars. Suggestive emVars (*P*_*emp*_<0.05) are shown in darker shades. **b**, Sharing of emVars across cell types and stimulation conditions. Spearman correlation of emVar effect sizes between conditions are shown for all 689 emVars (top) and for the subset reaching FDR<5% in both conditions (bottom).

**Extended Data Fig. 7.**
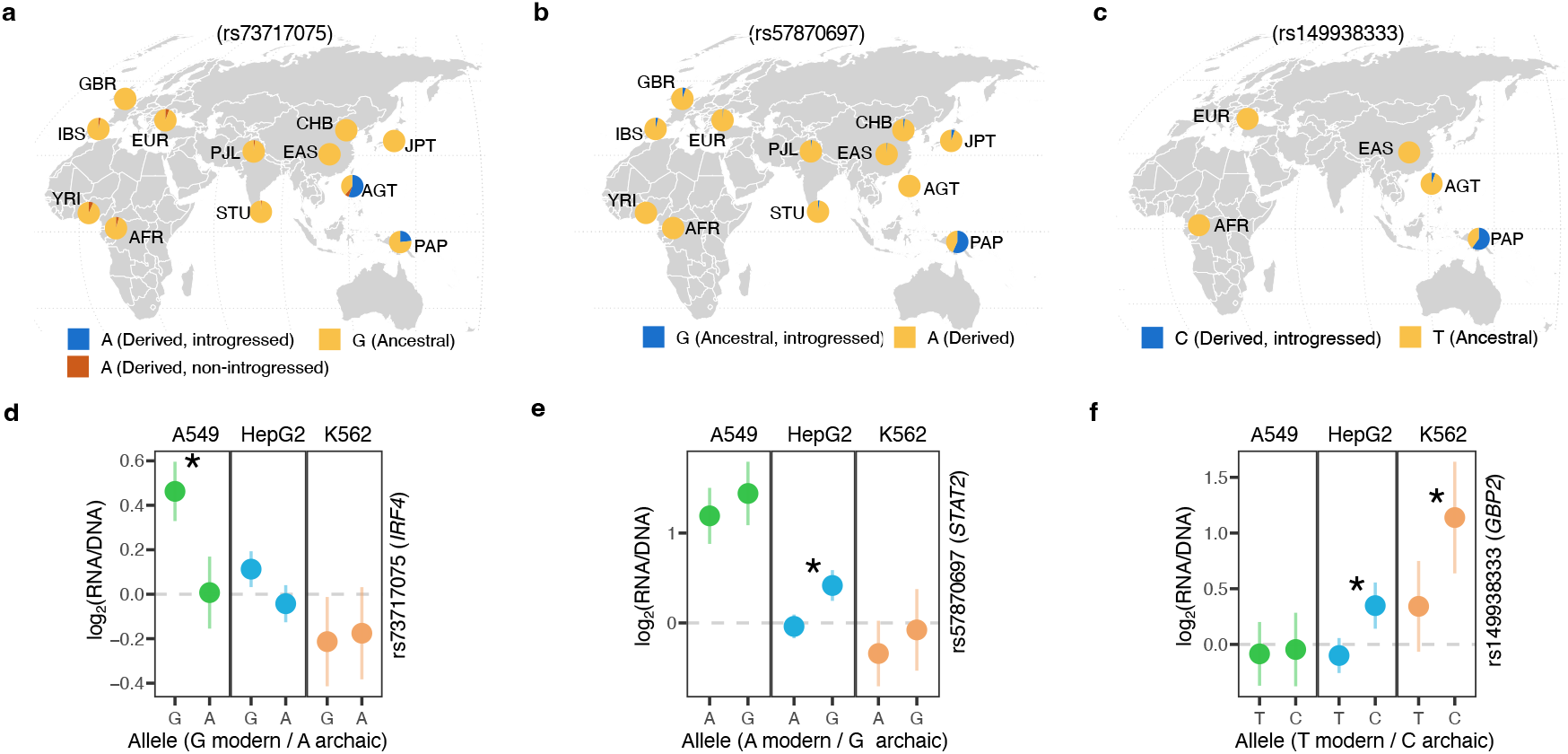
Examples of emVars in underrepresented populations. **a-c**, Worldwide allele frequencies of rs73717075-*IRF4* (**a**), rs57870697-*STAT2* (**b**) and rs149938333-*GBP2* (**c**) in the SGDP and 1KGP panels. **d-f**, Estimated cCRE activity of each allele of rs73717075 (**d**), rs57870697 (**e**) and rs149938333 (**f**) across the three tested cell types (error bars indicate 95% confidence intervals). Activity is shown for the basal (unstimulated) condition only.

**Extended Data Fig. 8.**
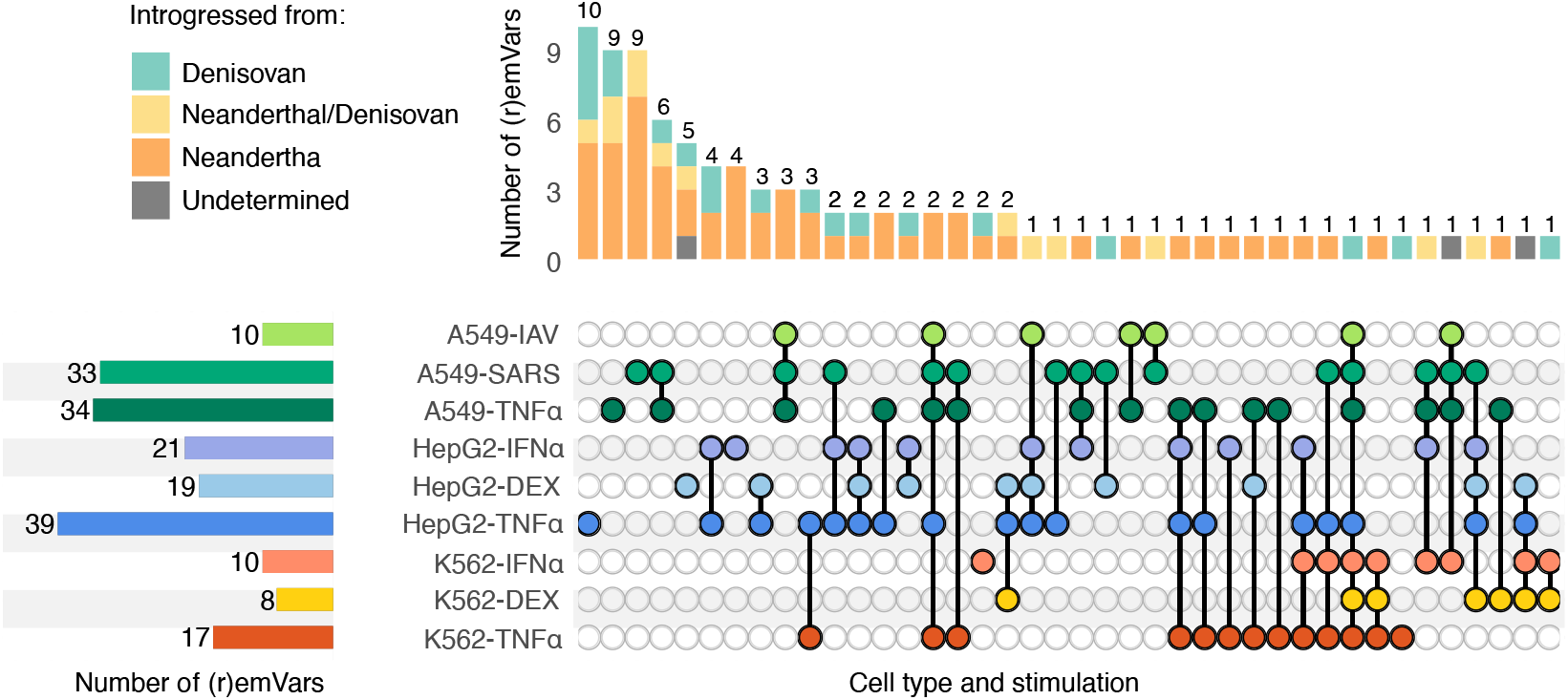
Sharing of responsive emVars across contexts (relaxed threshold). Upset plots show the frequency of responsive emVars across contexts (cell types × stimuli) and across specific combinations of contexts, using FDR_GxE_<5% for discovery and relaxed *p*-value threshold (*P*_*emp*_<0.05) for replication.

**Extended Data Fig. 9.**
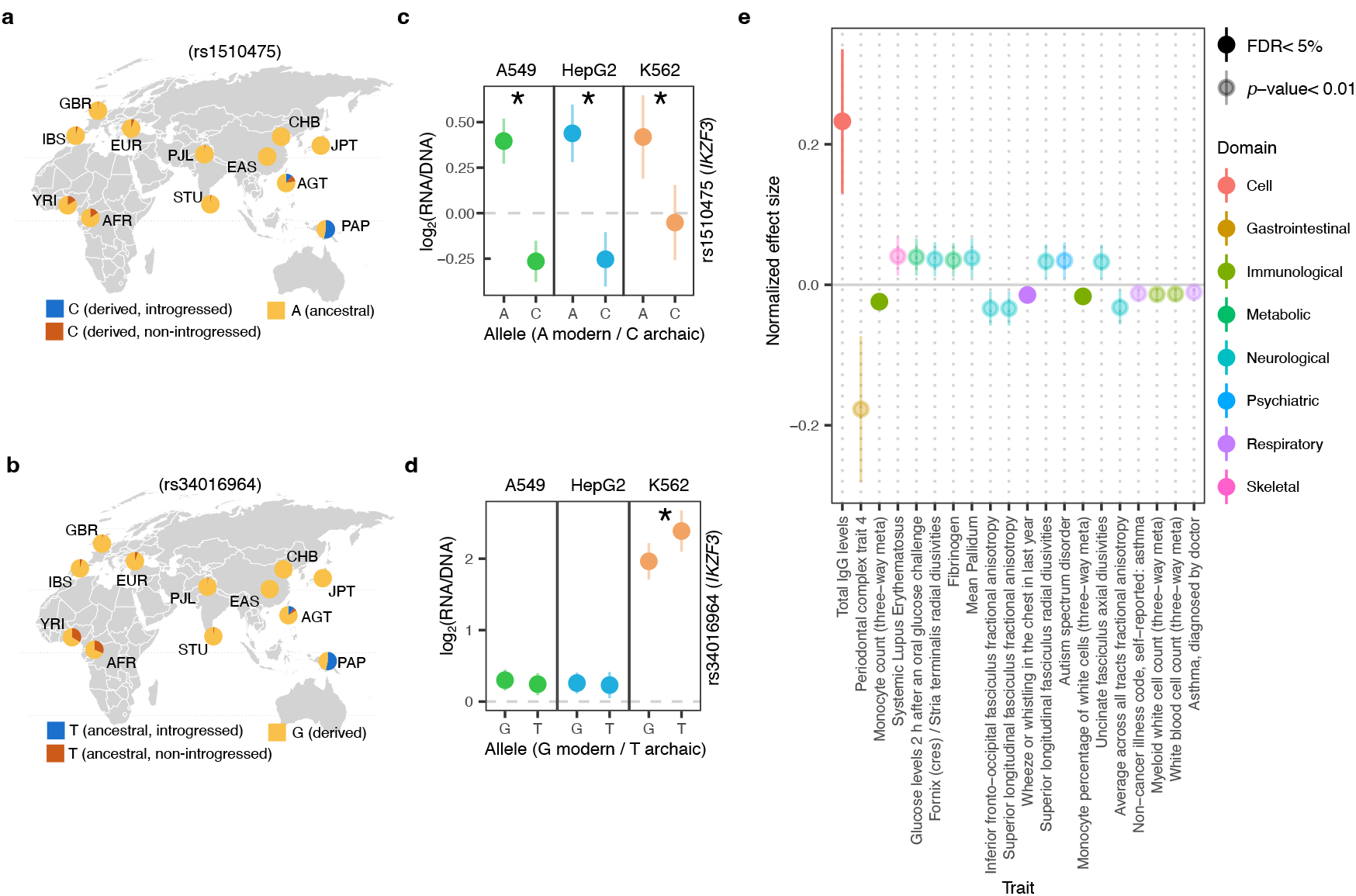
Examples of trait-associated emVars. Worldwide allele frequency (**a**,**b**) and estimated cCRE activity (**c**,**d**) for rs1510475-C and rs34016964-T. **e**, Phenome-wide association study of the rs9520848 variant. For each trait, the normalized effect size (in standard deviation units) and corresponding 95% confidence interval are shown, inferred from effect allele, *p*-value and sample size. Only the 20 traits with the largest absolute lower bound of the 95% confidence interval are displayed.

**Extended Data Fig. 10.**
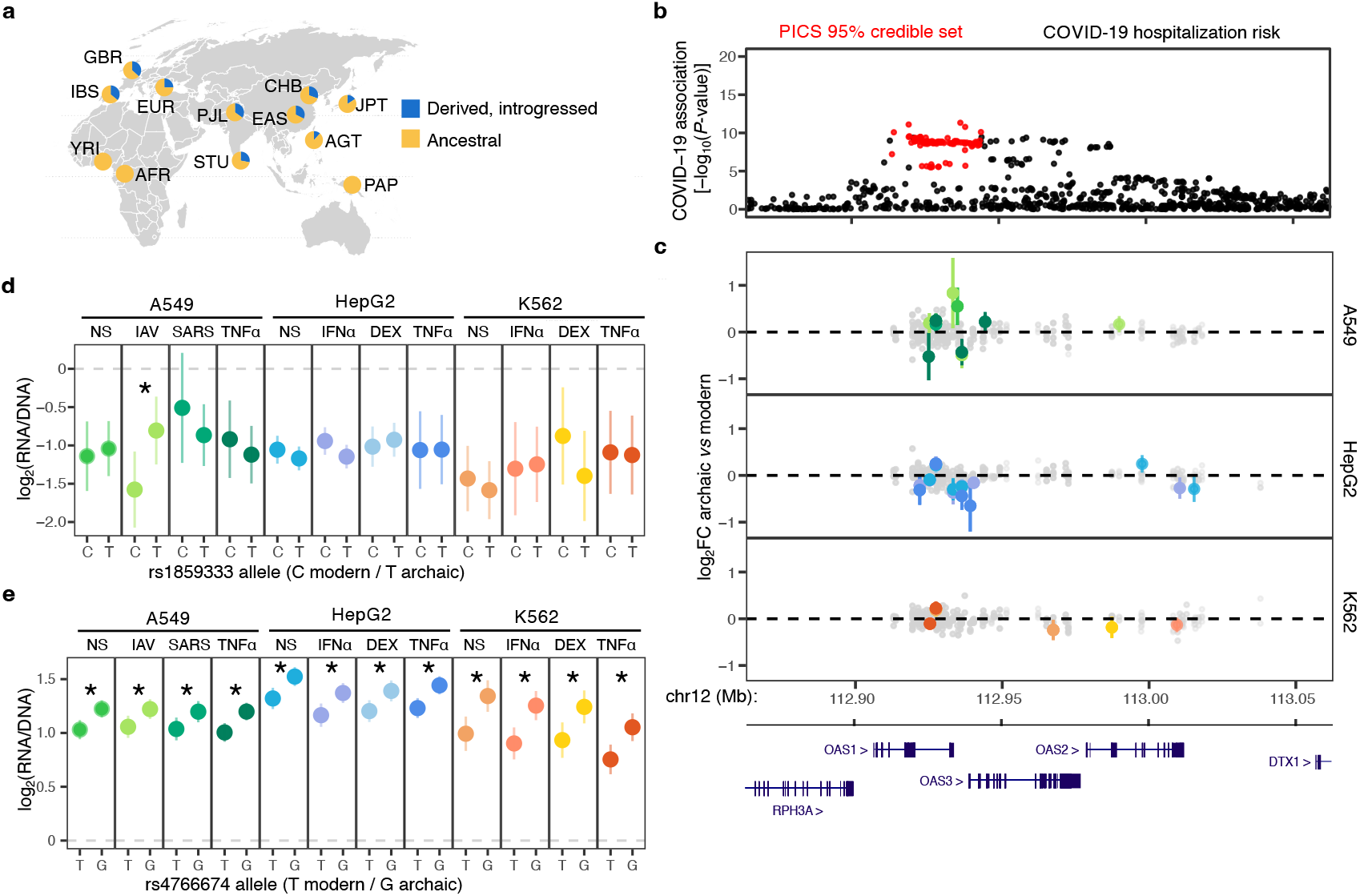
Regulatory effects of adaptively introgressed alleles at the *OAS1-3* COVID-19 risk locus. **a**, Worldwide allele frequency of introgressed alleles at the chromosome 12 *OAS1-3* COVID-19 locus (rs1859333 shown as a representative variant). **b**, LocusZoom plot of association with COVID-19 hospitalization risk (GCST90134600).Variants within the 95% credible set are highlighted in red. Protein-coding genes at the locus are shown below. **c**, MPRA-inferred regulatory effects. For each variant, the log_2_FC with 95% confidence intervals is shown across all cell types and stimulation conditions. Non-significant effects are shown in grey, whereas significant emVars (FDR<5%, |log_2_FC|>0.2) are colored by condition, as in (**d**,**e). d**,**e**, Estimated cCRE activity for each allele of rs185333 (**d**) and rs4766674 (**e**) across all tested conditions (error bars indicate 95% confidence intervals).Black stars denote emVars (FDR<5%).

