## Supplementary Notes1-6 and Figs 1-13 for "Immune context unmasks regulatory effects of Neanderthal and Denisovan introgression"

The **Supplementary Information** file contains Supplementary Notes 1-6, which include additional context, information on methodology, and additional results; Supplementary Figures 1-13; and Supplementary References.

### Supplementary Notes

#### Supplementary Note 1: Relevance of selected cell lines and technical considerations

Our MPRA experiments were performed across three different cell lines (A549, HepG2, and K562) to capture diverse aspects of human biology and immune response to infection. **A549** is a human alveolar epithelial cell line derived from a lung carcinoma, widely used to study respiratory viral infections including SARS-CoV-2 and influenza. In this study, we used A549 ACE2plus-C3 cells (ATCC, CRL-3560), which express ectopic ACE2 and TMPRSS2, rendering them highly permissive to SARS-COV-2 infection[1]. **HepG2** is a human hepatocellular carcinoma-derived liver cell line that retains many metabolic and immune functions of primary hepatocytes. Hepatocytes are key targets for hepatitis B and C viruses and contribute to the life cycle of other pathogens, such as *Plasmodium* species, which cause malaria[2]. They also play essential roles in drug and toxin metabolism and in the production of acute-phase proteins, including C-reactive Protein (CRP) and complement factors, which are critical for systemic immune responses[3]. **K562** is an immortalized myelogenous leukemia cell line derived from multipotent progenitor of erythroid and myeloid lineage[4]. K562 cells have been widely used as model of immune cells. The three cell lines were selected not only for their biological relevance but also for their robust growth and high infectivity by lentivirus, which is critical for MPRA experiments. Initial attempts using Calu-3 cells (lung epithelial proxy), Jurkat cells (T lymphocytes proxy), and THP-1 cells (myeloid lineage proxy) yielded insufficient infectivity, resulting in extremely low complexity of the MPRA library.

#### Supplementary Note 2: Choice of immune and infectious stimuli

To monitor the diversity of immune response across tissues, we stimulated cells with Interferon-alpha (IFN-α) to induce antiviral responses, Tumor Necrosis Factor-alpha (TNF-α) to trigger inflammation, or Dexamethasone (DEX) to suppress immune activity. These molecules were chosen for their well-characterized pro- or anti-inflammatory roles and the broad expression of their receptors[5-7]. IFN-α is a cytokine that activates the JAK-STAT signaling pathway, promoting expression of interferon-stimulated genes (ISGs) involved in antiviral defense and immune regulation[8]. TNF-α is a key pro-inflammatory cytokine that activates NF-κB, inducing genes involved in inflammation, apoptosis, and immunity; it is implicated in infections, auto-immune diseases, and cancers[9]. DEX is a synthetic glucocorticoid that interacts with the glucocorticoid receptor (GR, encoded by *NR3C1*) and regulates genes involved in metabolism, inflammation, and stress responses through glucocorticoid response elements (GREs)[10]. Clinically, DEX is used in patients presenting infectious diseases, for example to mitigate cytokine storms in severe COVID-19 patients[11]. Given the reported enrichment of adaptive introgression at loci encoding human proteins that interact with RNA viruses[12], in A549 cells we replaced IFN-α and DEX with live Influenza A Virus (H1N1) and SARS-Cov-2 to assess epithelial lung cell responses to actual exposure to RNA respiratory viruses. Both viruses can successfully infect the A549 cell line used in our MPRA experiment[1,13].

#### Supplementary Note 3: Adjustment and normalization of estimated activity

When quantifying cCRS activity, we observed that scrambled sequences showed a shift toward higher activities relative to test sequences ($\Delta\log_2$(RNA/DNA)>0.14, Wilcoxon rank sum $P<5.5\times10^{-6}$, Supplementary Fig. 2a). This shift was fully explained by the high GC-content of scrambled sequences, which were derived from GC-rich promoter regions (mean GC content: 59% vs. 44% for test sequences, Wilcoxon rank sum $P<1.6\times10^{-39}$, Supplementary Fig. 2b). Indeed, among scrambled sequences, cCRS activity increased on average by 1.2% for each additional percent of G or C nucleotides (Student $P<2.7\times10^{-19}$, Supplementary Fig. 2c). Adjusting activity for GC-content not only improved the comparability of activity levels between scrambled and test sequences (Fig. 1d), but also revealed enrichment of polycomb-repressed regions among downregulating cCREs (Extended Data Fig. 3b) and increased concordance in activity between replicates (Supplementary Fig. 2d). GC-correction also increased by 36-72% the number of sequences considered as active in the K562 cell line, where the signal-to-noise ratio was generally lower, while having limited effects in other cell lines. In addition, because the distribution of estimated cCRS activities varied across cell types in both observed and permuted data (Supplementary Fig. 2e), we scaled activities across all cCRS to achieve identical means and variances across conditions. The resulting GC-corrected, normalized activities were used to define active CREs and for visualization of emVars and responsive cCREs. Differential activity analyses (e.g. responsive cCREs and emVars), however, were based on raw UMI counts and were therefore unaffected by these normalization procedures.

#### Supplementary Note 4: Global transcriptional shutdown decreases cCRE activity

To test whether the decreased activity of both upregulating and downregulating cCREs observed upon viral stimulation could be explained by a global transcriptional shutdown in a subset of infected cells, we simulated cCRS activity in cells that were either resting or stimulated with a virus inducing shutdown of RNA production upon infection. Specifically, we simulated $n_b$=1000 barcoded cCRS integrated across $p$=1000 cells (assuming a single barcode per cCRS for simplicity), with an average of $n_i$=200 integrations per cell (DNA UMIs). We assumed that, upon viral stimulation, only a fraction $\pi$=20% of cells remained healthy (bystander cells), whereas the remainder were infected. We further assumed that the total number of sequenced transcripts (RNA UMIs) in cell $c$, denoted $U_c$, were normally distributed with mean $\mu$ and standard deviation $\sigma$ that differed between healthy and infected cells. We set $\mu_h = 8000, \sigma_h = 10$ for healthy cells and $\mu_i = 20, \sigma_i = 3$ for infected cells. Finally, we sampled the fixed transcriptional activity $\alpha_i$ of each cCRS $i$, which determines the normalized RNA/DNA ratio of the associated barcode, from a log-normal distribution (mean 0, SD 1). For each cell $c$, we then sampled $U_c$ transcripts, assigning barcode $i$ with probability $p_{ic} = \frac{\alpha_i}{\sum_{i \in \mathcal{B}_c} \alpha_i}$, where $\mathcal{B}_c$ is the set of barcodes associated with cCRS integrated in cell $c$. For each condition (resting or viral stimulated cells) and cCRS $i$, we computed the total number of cells in which cCRS $i$ was integrated (DNA UMIs, denoted $d_i$) and the total number of transcripts it produced (RNA UMIs, denoted $r_i$). In each condition, the activity of cCRS $i$ was estimated as $\hat{\alpha}_i = \frac{r_i / \sum_i r_i}{d_i / \sum_i d_i}$. Simulations (Supplementary Fig. 11) showed that, in the viral condition, where infected cells underwent global transcriptional shutdown without changes in cCRE activity, estimated transcriptional activity was reduced of both upregulating and downregulating cCREs (ie. $\hat{\alpha}_{STIM} < \hat{\alpha}_{NS}$ for upregulating cCREs and $\hat{\alpha}_{STIM} > \hat{\alpha}_{NS}$ for downregulating cCREs), recapitulating the patterns observed experimentally following IAV and SARS-CoV-2 stimulation.

#### Supplementary Note 5: Comparison with eQTL and other emVar studies

To evaluate the validity of identified emVars, we first intersected them with eQTL databases[14]. Of the 342 emVars identified at basal state, 34% overlapped fine-mapped credible sets from the eQTL catalog, with this proportion ranging from 33 to 37% across conditions. Among all emVars overlapping an eQTL, 71% showed concordant allelic effects in at least one tissue, consistent with previous estimates[15-17]. However, 29% of emVars exhibited effects opposite to the eQTL direction, and 32% showed context-dependent associations with expression, correlating with both increased or decreased gene expression depending on the target gene or tissue. These observations suggest that sequence context, feedback loops, and additive effects of linked genetic variants play a critical role in shaping the regulatory impact of introgressed alleles.

We next compared allelic effects measured in resting cell lines with those identified in a large-scale plasmid-based MPRA study[18]. Among 160 variants shared with our study, 28 were detected as emVars in either ref.[18] or our study, with only 3 being significant in both (OR=3.1, Fisher's exact *P*=0.12). Nevertheless, 87% of detected emVars showed consistent effect directions across studies (95% CI: [0.70-0.96]), indicating good replicability despite limited power for detection (Spearman ρ= 0.51, *P*=0.003, Supplementary Fig. 12a). Notably, emVars reported in ref.[18] showed consistently lower effect sizes upon replication (Wilcoxon $P<9.0\times10^{-8}$, Supplementary Fig. 12b), whereas emVars from our study maintained similar effect sizes (Wilcoxon *P*=0.33), consistent with the wider dynamic range of plasmid-based MPRA relative to lentiviral-based approaches[19].

#### Supplementary Note 6: Power calculations

To evaluate our ability to detect regulatory variants across cell types and stimulations, we simulated regulatory variants across a wide range of effect sizes by randomly pairing cCRS and treating them as pseudo-alleles of a shared variant (Supplementary Fig. 13a). We then resampled DNA and RNA UMIs for each cCRS using a binomial bootstrap strategy: given the vectors $d^o$ and $r^o$, representing observed DNA and RNA counts for all barcodes and replicates, we resampled UMIs vectors for DNA, $d^s$, and RNA, $r^s$, by sampling counts of barcode $i$ in replicate $j$ from a binomial distribution, adding an offset of 1 to maintain the total number of detected barcodes constant:

$$d_{ij}^s \sim 1 + \mathrm{Bin}(\,p_{ij}^d, N^d)$$

and

$$r_{ij}^s \sim 1 + \mathrm{Bin}(\,p_{ij}^r, N^r)$$

where $p_{ij}^d = \frac{d_{ij}^o}{\sum_{ij} d_{ij}^o}$ and $p_{ij}^r = \frac{r_{ij}^o}{\sum_{ij} r_{ij}^o}$ are the fractions of UMIs associated with barcode $i$ and replicate $j$, in all DNA and RNA barcodes, respectively; and $N^d = \sum_{ij}(d_{ij}^o - 1)$, and $N^r = \sum_{ij}\left(r_{ij}^o - 1\right)$ represent the total number of informative UMIs (excluding the first UMI that ensures barcode detection) in DNA and RNA. The resampled vectors $d^s$ and $r^s$ were used to estimate cCRS activity while mimicking random capture of barcodes during sequencing, allowing assessment of our ability to detect true differential expression (under the alternative hypothesis $H_1$: $|log_2FC|>0$, Supplementary Fig. 13b). The observed DNA and DNA counts, $d^o$ and $r^o$, serve to estimate the true activity of each cCRS and derive the true effect size of the pseudo-variant.

Power was estimated either across conditions by binning pseudo-variants according to total number of barcodes × replicates (Supplementary Fig. 13c) or within condition by binning according to effect size alone to provide the average power across all cCRS in a given condition, which depends on both the observed number of replicates and the distribution of barcodes per cCRS (Supplementary Fig. 13d). Two *p*-value thresholds were considered: a 5% FDR (as used in our study) and a 5% empirical *p*-value threshold for cross-tissue replication. Our analysis shows that ~50 observations per allele (barcodes × replicates) are sufficient to detect strong effects ($|log_2FC| >0.5$) with ~70% power, whereas ≥400 observations are required to detect weaker effects ($|log_2FC| >0.2$) with ~84 % power. Reducing the *p*-value threshold for replication allows recovery of weak effects from much fewer observations (>80% power for emVars with $|log_2FC|>0.2$ for 150 observations only).

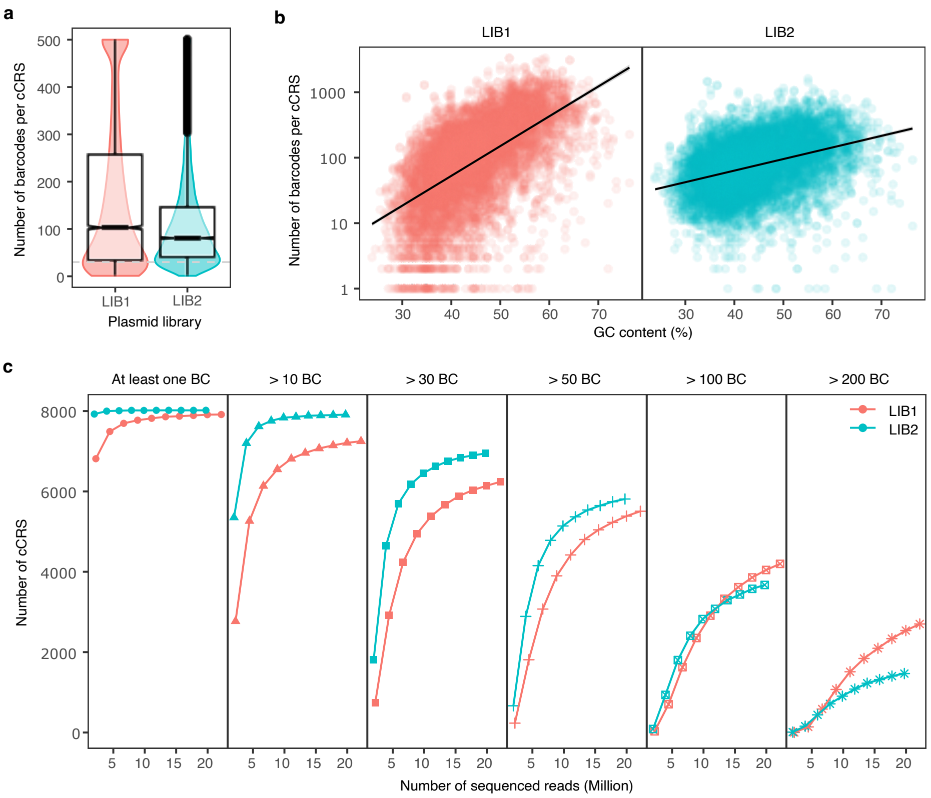


**Supplementary Fig. 1 | Barcode-oligo association results.** Two plasmid libraries were generated. For the second library, PCR enzyme conditions were optimized to reduce GC-contents bias, and the target number of barcodes per cCRS was reduced from 200 to 100 to optimize sequencing cost. **a**, Distribution of the number of barcodes per cCRS across plasmid libraries. **b,** Number of barcodes per cCRS strongly correlates with GC content. **c**, Saturation analysis showing the number of cCRS recovered in each library as a function of the minimal number of barcodes required.

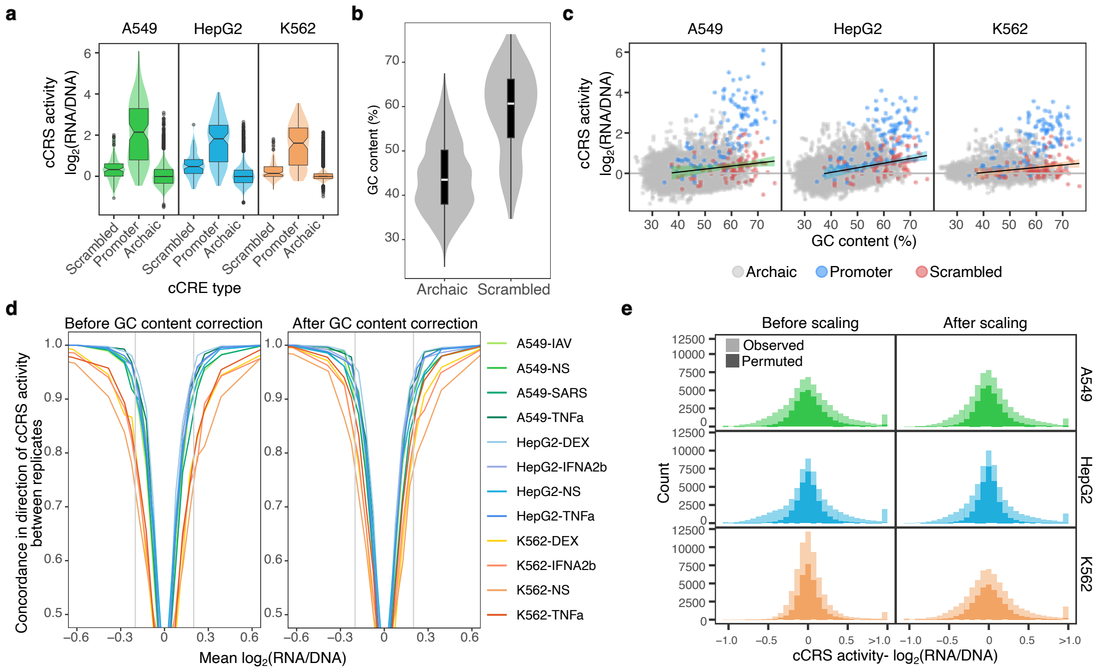


**Supplementary Fig. 2 | Impact of GC content on MPRA activity. a**. Violin plots showing the distribution of raw transcriptional activity for random sequences (scrambled controls), promoters (positive controls), and tested introgressed sequences across all three cell types (unstimulated condition). **b**. Distribution of GC content among introgressed and scrambled cCRS. **c**. Scatter plots showing uncorrected activity distributions (1 dot = 1 cCRS) in the unstimulated condition across all three cell lines. Positive controls are shown in blue, and their scrambled counterparts in red. Regression lines are fitted using scrambled sequences only. **d**. Mean concordance in direction of cCRS activity across replicates, reported as a function of the mean log2(RNA/DNA) ratio, separately for each condition, with or without GC correction. **e**. Distribution of observed and permuted cCRS activities in each cell type (unstimulated condition), before and after scaling. Figure associated with Supplementary Note 3.

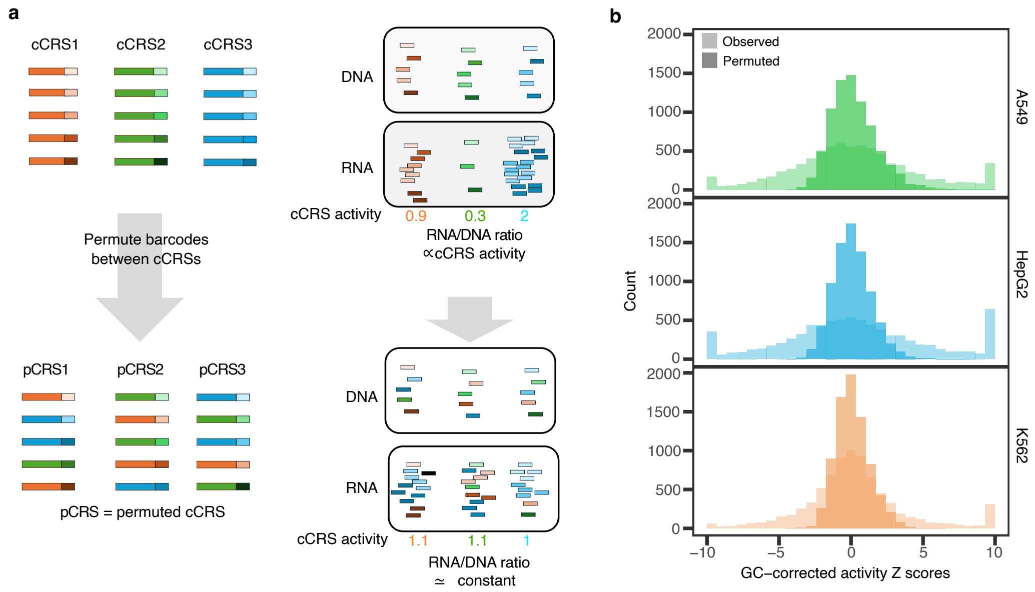


**Supplementary Fig. 3 | Identification of active cCRS through barcode permutation**. **a**, Schematic of the permutation approach. Each cis-regulatory sequence (cCRS, numbered 1-3 and colored in orange, blue, and green) is associated with distinct barcodes, each represented by different shades of orange, blue, and green. In observed data, RNA/DNA ratios of barcodes reflect cCRS activity. Permuting barcodes across sequences generates pseudo-cCRS (pCRS) with matched barcode composition but no regulatory signal, yielding expression levels comparable to inactive sequences. **b**, Distribution of activity Z-scores for observed and pCRS across three cell lines (unstimulated condition).

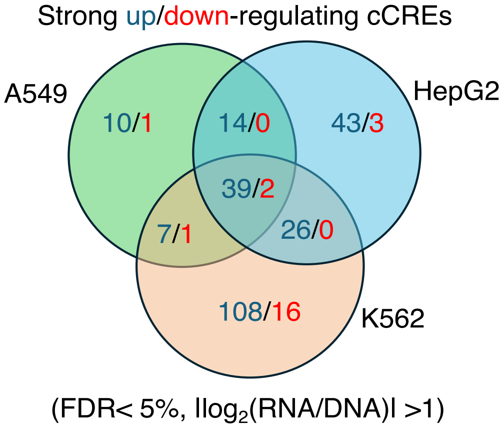


**Supplementary Fig. 4 | Strongly active cCREs are highly cell type-specific.** Overlap of strongly active cCREs across cell types (FDR<5%, $|\log_2(\text{RNA/DNA})|>1$, unstimulated condition). Numbers are shown separately for upregulating (blue) and downregulating (red) cCREs.

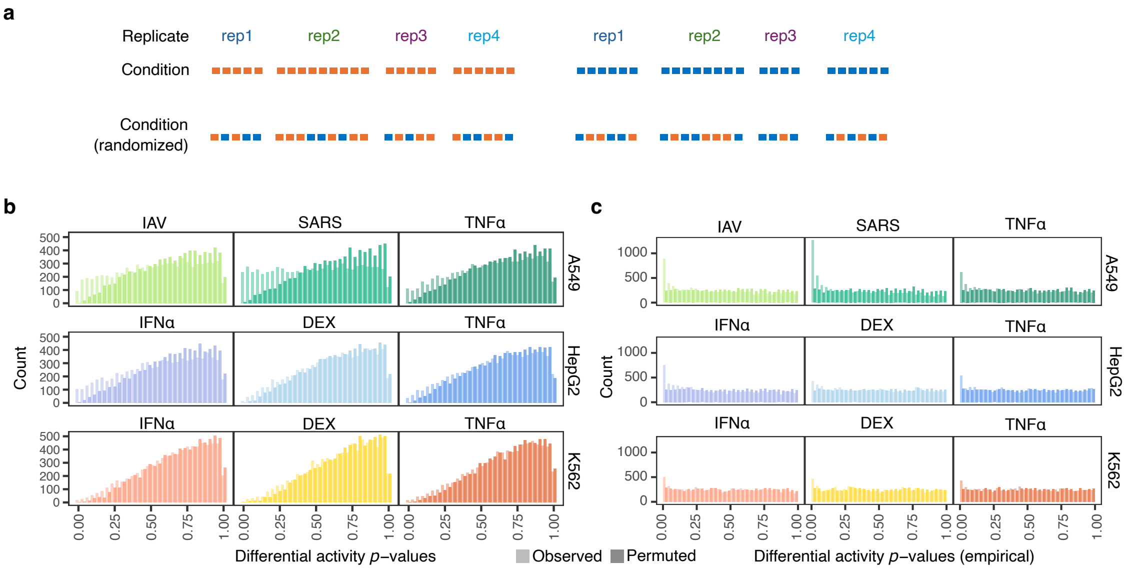


**Supplementary Fig. 5 | Permutation strategy for differential activity testing across samples (responsive cCREs). a,** Schematic of the permutation approach. In our design, each replicate is associated with a single condition (shown in orange and blue), with multiples barcodes providing independent measurements of cCRS activity in that condition. Random Reassigning of each measurement (barcode × replicate) at random across conditions generates artificial cCRS with no true differential activity between conditions, forming a null distribution. **b**, Distributions of differential activity *p*-values obtained from *MPRAnalyze* across all stimulations, with and without permutation. Permuted *p*-values shift toward 1 indicates a bias toward the null hypothesis. **c,** Empirical *p*-values derived from permutations show a flat distribution under the null and enrichment toward 0 in observed data.

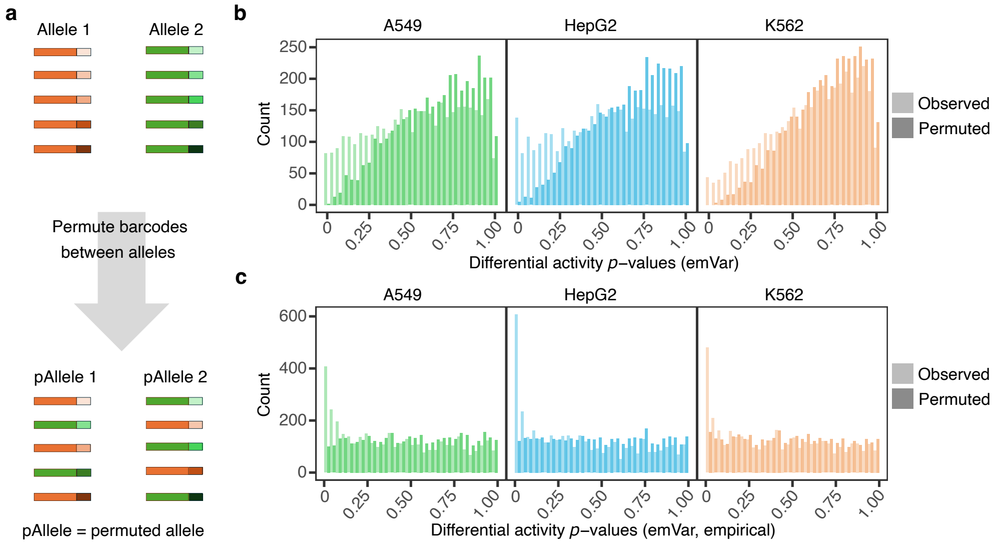


**Supplementary Fig. 6 | Permutation strategy for emVar testing. a,** Schematic of the permutation approach. Each SNP is represented by two alleles (orange and green), each associated with distinct barcodes (different shades of orange and green). Random reassignment of barcodes between alleles generates artificial SNPs with no allelic effect, defining a null distribution for emVar testing. **b,** Distribution of differential activity *p*-values from *MPRAnalyze* across all three cell lines, with and without permutation. Permuted *p*-values shift toward 1 indicates a bias toward the null hypothesis. **c,** Empirical *p*-values from permutations show a flat distribution under the null and enrichment toward 0 in observed data.

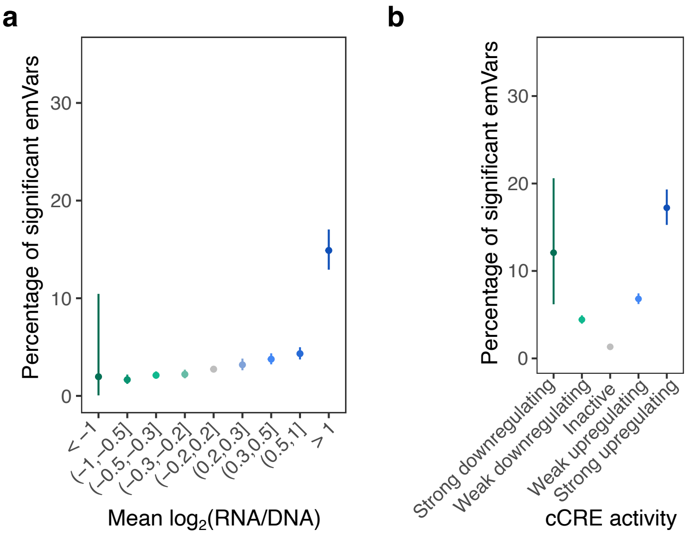


**Supplementary Fig. 7 | Enrichment of EmVars in active cCREs.** Proportion of significant emVars (FDR<5%, $|log_2FC|>0.2$) as a function of cCRE activity, with 95% confidence intervals **a**, Binned by estimated cCRE activity (irrespective of significance) and **b,** Grouped by class of cCRE activity.

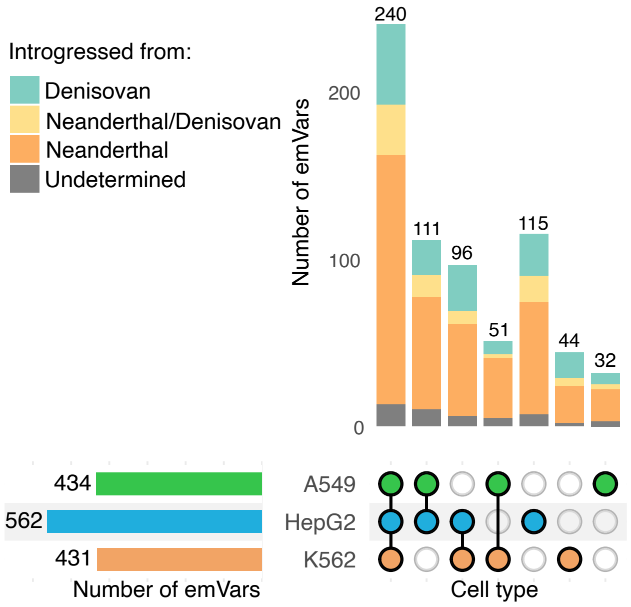


**Supplementary Fig. 8 | Sharing of emVars across cell types (relaxed threshold).** Upset plots showing the number of emVars detected across combinations of cell types, using FDR<5% for discovery and relaxed *p*-value threshold ($P_{emp}$<0.05) for replication.

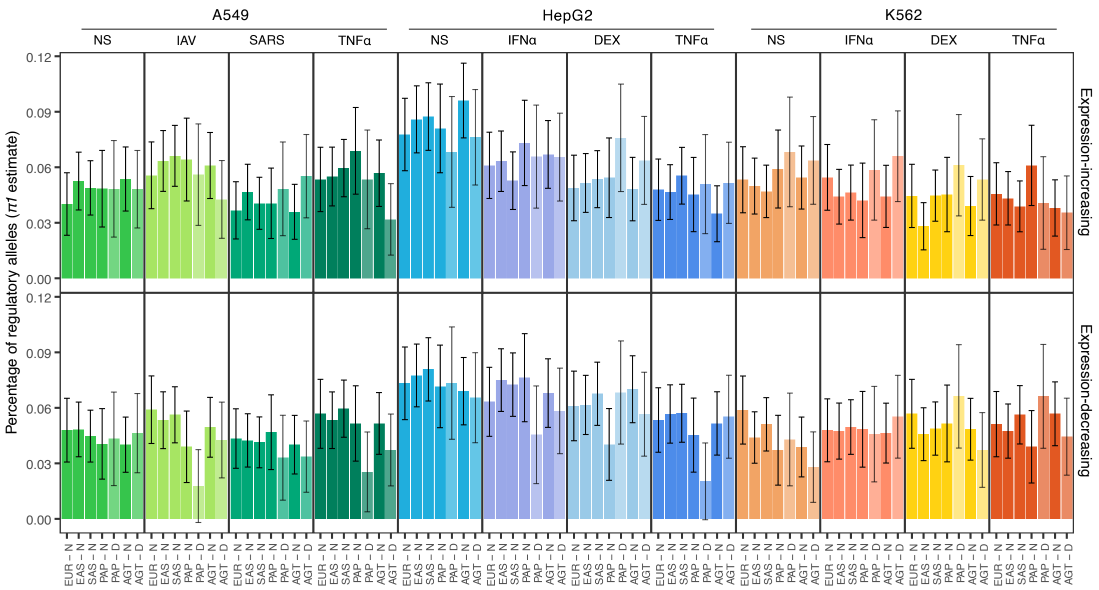


**Supplementary Fig. 9 | Breakdown of regulatory alleles by population, archaic source, and effect direction.** Estimated proportion ($\pi_1$) of Neanderthal (N) and Denisovan (D) alleles that increase of decrease reporter gene expression across populations and conditions. Bars indicate 95% bootstrap confidence intervals (extended from Fig. 5b across all conditions).

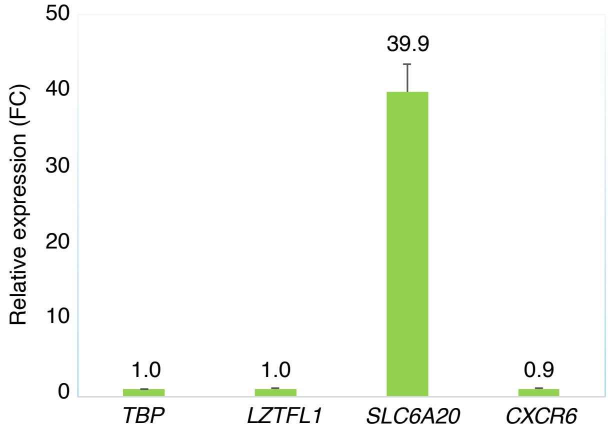


**Supplementary Fig. 10 | Expression changes of *LZFTL1*, *SLC6A20*, and *CXCR6* genes upon CRISPR activation.** qPCR-based fold changes following CRISPR activation of the rs11713054-associated enhancer. Error bars indicate 95% confidence intervals. Relative gene expression levels were normalized to housekeeping gene *TBP*.

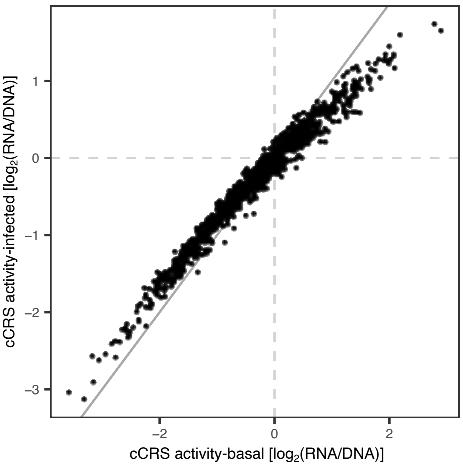


**Supplementary Fig. 11 | Effect of a global transcriptional shutdown on estimated activity.** Simulated global transcriptional shutdown in infected cells without changes in cCRE activity results in reduced estimated activity for both up- and downregulating cCREs, similar to that observed in the viral stimulation conditions. Figure associated with Supplementary Note 4.

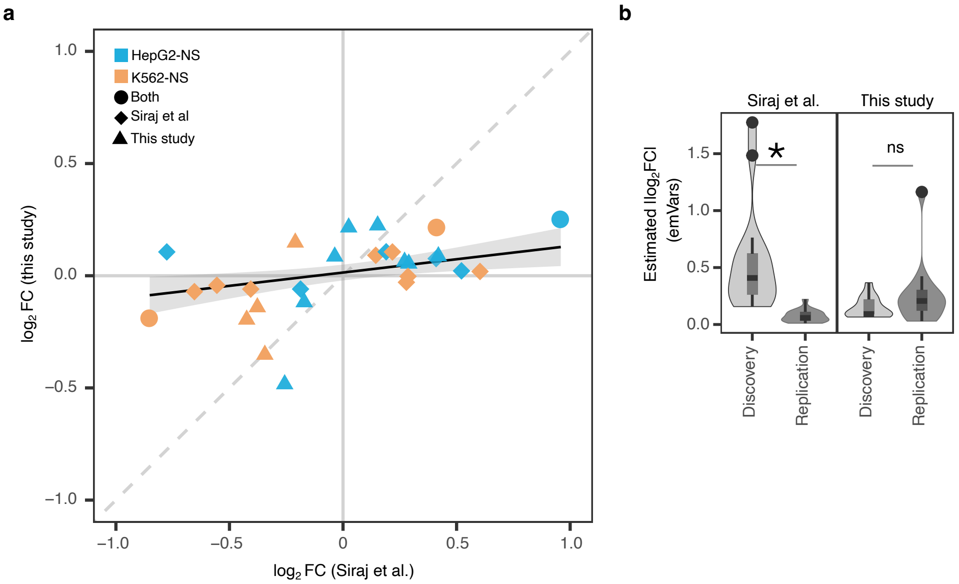


**Supplementary Fig. 12 | Comparison of emVars with plasmid-based assay**. Comparison of emVars identified in our study (resting K562 and HepG2 cells) with results from Siraj *et al.*[18] for 160 shared SNPs. **a**, Scatter plot of effect sizes across studies. Only SNPs identified as emVars in at least one study are shown. Shape indicates study of emVar discovery; color indicates cell type. **b**, Distribution of effect sizes for emVars reported in Siraj et al (left) and in this study (right). * Wilcoxon ram-sum test $P<10^{-3}$; ns: not significant ($P>0.05$). Figure associated with Supplementary Note 5.

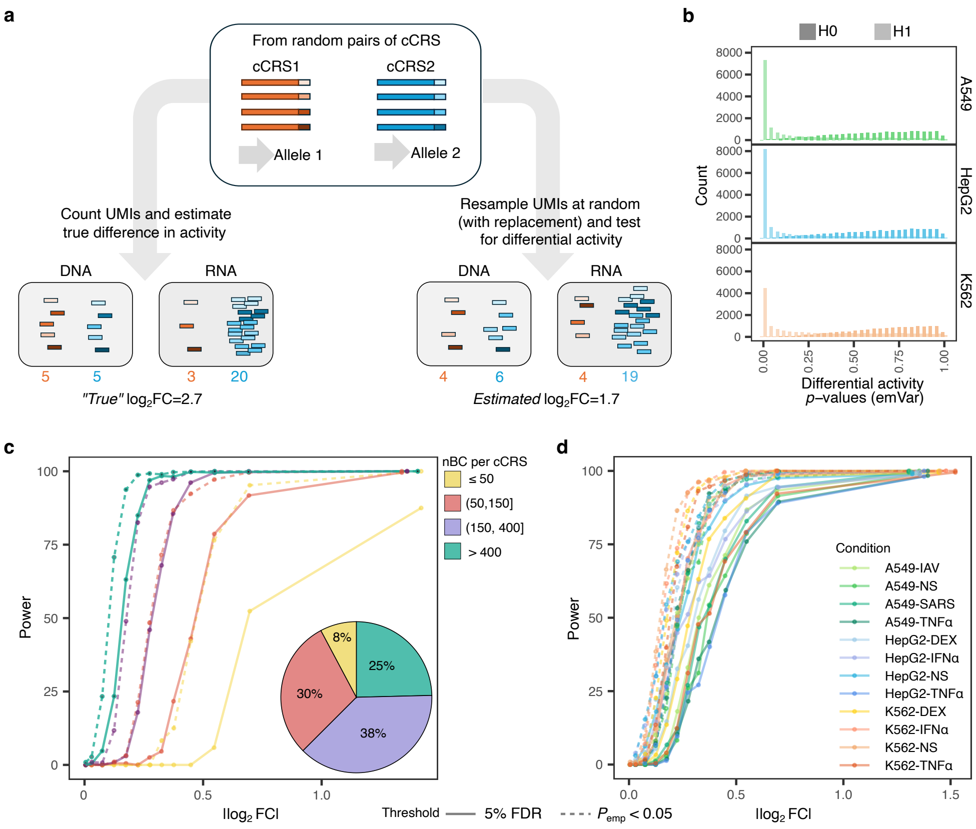


**Supplementary Fig. 13** | Power analysis for emVar detection. **a**, Power estimation framework based on synthetic emVars generated from random cCRS pairs. We compute the difference in activity between the two cCRS, before resampling DNA and RNA UMIs at random to mimic the capture of DNA and RNA barcodes during sequencing. We use these resampled UMIs to test for differential activity and assess our ability to detect emVars across a wide range of effect sizes. **b,** Distribution of differential activity *p*-values from synthetic emVars simulated under $H_1$ ($|\log_2 FC > 0|$) across all three cell types (unstimulated condition). The permutation scheme described previously (see Supplementary Fig. 6) is used for the null hypothesis ($H_0$: $|\log_2 FC| = 0$). **c,** Power for emVar discovery increases drastically with effect size ($\log_2 FC$) and with the number of observations (barcodes × replicates) per allele. Pie chart shows the distribution among tested SNPs of the number of observations per allele. **d**. Estimated power across conditions based on observed barcode distributions. In **c**,**d**. solid lines indicate power for initial discovery (FDR<5%, and $|\log_2 FC| > 0.2$), and dashed lines indicates power for replication (empirical $P < 0.05$). Figure associated with Supplementary Note 6.
